# Comprehensive Profiling of Plasma Exosomes Using Data-Independent Acquisitions – New Tools for Aging Cohort Studies

**DOI:** 10.1101/2021.02.27.433188

**Authors:** Sandip K. Patel, Roland Bruderer, Nathan Basisty, Joanna Bons, Pierre-Yves Desprez, Francesco Neri, Lukas Reiter, Judith Campisi, Birgit Schilling

## Abstract

Aging is a complex biological process associated with progressive loss of physiological function and susceptibility to several diseases, such as cancer and neurodegeneration. Exosomes are involved in many cellular signaling pathways, and their cargo may serve as promising disease or aging biomarkers. These membrane-bound extracellular vesicles facilitate the transport of intracellular contents to proximal and distal cells in the body. Here, we investigated two omics approaches for exosome analysis. To overcome the challenges of plasma exosome contamination with abundant soluble plasma proteins, we developed a high-throughput method to isolate highly purified exosomes from human plasma by sequential size-exclusion chromatography and ultrafiltration. First, we used data-dependent acquisitions from offline high-pH reversed-phase fractions of exosome lysate to generate a deep spectral library comprising ∼2,300 exosome proteins. Second, in a pilot aging study, we used comprehensive data-independent acquisitions to compare plasma exosomes from young (20–26 yrs) and old (60–66 yrs) individuals. We quantified 1,318 exosome proteins, and levels of 144 proteins were significantly different in young and old plasma groups (Q<0.05 and >1.5-fold change). We also analyzed exosome miRNA cargo and detected 331 miRNAs. Levels of several were significantly different in young and old individuals. In addition, 88 and 17 miRNAs were unique to old and young individuals, respectively. Plasma exosome biomarkers have great potential for translational studies investigating biomarkers of aging and age-related diseases and to monitor therapeutic aging interventions.

## Introduction

Aging demographics have changed remarkably over the last century, leading to older populations and subsequent stresses on the healthcare system and society. By 2030, ∼20% of North Americans are predicted to be 65 years and older. Aging is strongly correlated with cardiovascular diseases (CVD), cancer, osteoarthritis, cognitive decline and other disorders.^1, 2^ Thus, efficient predictive Markers of Aging are an urgent need.

To develop aging biomarkers, one strategy involves senescence-derived biomarkers for aging.^3^ Senescence burden increases with age, ^4^,^5^ and many age-related conditions are correlated to senescence burden.^6^ Interestingly, the senescence-associated secretory phenotype (SASP)^7^ provides an excellent pool of protein candidates that may develop into biomarkers of aging.^8^ Thus, we recently reported a SASP Atlas resource (SASPAtlas.com) that characterized both the heterogeneous *soluble SASP*, consisting of secreted soluble proteins, and proteins identified in released exosomes or also referred to as *exosome SASP*. ^8^

Aging can be driven or mitigated by changes in circulating factors,^3, 8-10^ including circulating exosomes ^11, 12^ For example, D’Anca et al. recently reported that the unique exosome cargo composition related to specific physiological states of cells could serve as the plasma biomarker for aging and age-related diseases.^13^ Therefore, identifying the cargo of exosomes in plasma and other biofluids may contribute to identifying novel aging markers and pro- and anti-aging factors. Moreover, micro (mi)RNAs packaged inside exosomes circulate as exosome cargo in biofluids, and hence, their analysis could complement the proteomics analysis.^14^ Exosome cargo molecules may serve as markers for disease diagnosis and prognosis.^15-17^ Besides, exosomes are present in many biological fluids, such as urine,^18, 19^ saliva,^20^ semen,^21^ breast milk,^22^ plasma,^23^ amniotic fluid,^24^ serum,^25^ malignant ascites,^25^ pleural effusion,^26^ and various cell types that come in contact with fluids.^27, 28^ Thus, the abundance and composition of exosome miRNAs differ in diseased and healthy individuals, and several studies profiled exosomal miRNAs that may have applications for clinical diagnosis.^29-31^

However, the isolation and analysis of circulating exosomes or other EVs in biological fluids is challenging. The dynamic range of proteins in most biofluids, especially plasma makes it challenging to identify and quantify proteins, particularly low-abundance proteins, using modern mass spectrometry-based proteomics. Recently, exosomes gained enormous interest as sources of disease biomarkers: they are abundant in virtually all biofluids, have significant roles in cell-cell communication, and largely circumvent issues with the protein dynamic range. The major obstacle is efficient exosome isolation providing a high degree of enrichment and ‘exosome purity’,^32-34^ free of contaminating proteins, such as albumin, commonly found in biofluids. Current isolation methods rely on size differences among EVs or on target-specific surface markers.^35, 36^ Commonly used methods include ultracentrifugation (the current ‘gold standard’),^36, 37^ precipitation, filtration, chromatography, and immunoaffinity-based approaches. Commercial exosome isolation kits may lead to variable exosome purity,^38-41^ low yields, or may use reagents, such as PEG-based enrichments that are not compatible with downstream mass spectrometry-based proteomic applications. Size exclusion chromatography (SEC) is often employed prior to centrifugation or filtration to enhance exosome purity.^42-44^

In this study, we combined efficient exosome isolation with modern, comprehensive analytical approaches, such as data-independent acquisition-mass spectrometry (DIA-MS). DIA is a comprehensive and quantitative approach to proteomics analysis that offers significant advantages over the traditional data-dependent acquisition (DDA) approaches. DIA is highly reproducible, as it fragments all precursor ions instead of stochastically sampling the most abundant precursor ions (as in DDA), hence DIA is considered unbiased and provides flexibility to re-query and re-analyze the data post-acquisition.^45^ DIA offers rapid identification of thousands of proteins with accuracy and precision due to the continuous collection of fragment-ion spectra with fewer interferences over a preselected m/z range of precursor ions, even when applying shorter gradients.^46^ DIA workflows often include the initial generation of spectral libraries using traditional discovery methods, such as DDA coupled to offline HPLC fractionation, to create comprehensive spectral libraries with matching retention times to identify peptides from subsequent DIA analyses. However, the latter approach increases instrument acquisition time, and especially for small experiments, presents a significant cost overhead, unless an existing spectral library is available from previous work within the laboratory or from published work (e.g., pan-human library).^47^ Multiple library-free methods also exist for DIA analysis, such as DIA-Umpire,^48^ and directDIA.^49^ These workflows and algorithms to build spectral libraries directly from DIA acquisitions are rapidly improving.

In summary, we report an optimized DIA analysis method for human plasma exosomes using both library-coupled and library-free approaches (Fig. 1) investigating a small Aging Pilot Study (Table S1). We generated a deep plasma exosome spectral library (2323 proteins) using DDA acquisitions. We demonstrate that our plasma exosome protein library is highly enriched for known exosome protein annotation described in the ExoCarta database^50^ that provides a comprehensive web-based collection of exosome proteins, RNA, and lipids from various organisms. Finally, we introduce an alternative quantitative plasma exosome DIA-only workflow in which DIA acquisitions were searched directly in Spectronaut (Biognosys: directDIA) without a spectral library. The flexibility of combining different mass spectrometric scan types and prior knowledge will provide excellent future tools for biomarker research and comprehensive analysis of large human cohorts.

**Fig. 1.**
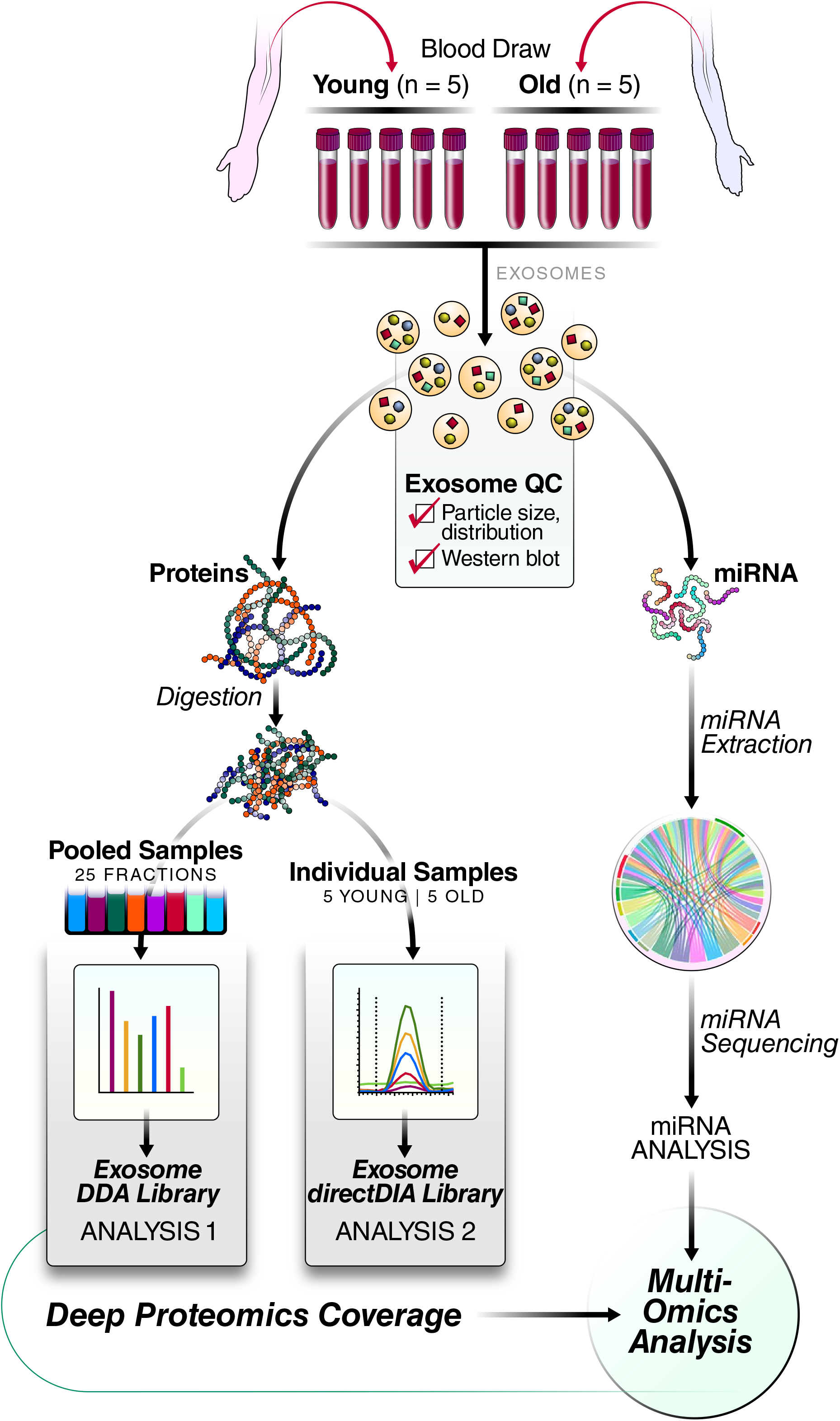
Multi-omics workflow for isolation and analysis of human plasma exosome proteins and miRNA. Plasma exosomes from young (n=5) and old (n=5) donor cohorts were extracted and assessed by particle-size distribution for size and intactness and western blots for their protein yields, purity, and potential contaminations with the plasma proteins. Exosome proteins were digested and subjected to DIA mass spectrometric analysis with two in-house-generated spectral libraries, using either i) a deep DDA library that was generated from 25 offline HPRP fractions (from pooled samples), or ii) a directDIA library generated from the 10 individual plasma DIA acquisitions. Exosome miRNA was extracted and analyzed in parallel.

## Results

### Plasma exosome extraction and characterization

To optimize the protocol for efficient plasma exosome enrichment achieving high exosome purity levels (with low soluble plasma protein contaminations) for downstream omics analysis, we performed plasma exosome extraction by four approaches (workflow shown in Fig. S1): i) ultracentrifugation (UC), ii) size-exclusion chromatography (SEC), as well as combinations iii) SEC/UC and iv) SEC and ultrafiltration SEC/UF.

UC is typically considered the “gold standard” for isolating exosomes from biofluids. Hence, we used UC as a benchmark of the exosome quality. In tissue-culture experiments with IMR90 cells (Fig. S2) and parallel plasma (Fig. 2) exosome experiments, UC enriched the exosomes as confirmed by the presence of exosome-specific markers, tumor susceptibility gene 101 (TSG101) and CD9, compared with soluble protein fractions or non-enriched fractions. For tissue-culture-derived exosome enrichments, soluble protein contamination, as assessed by alpha-2-macroglobulin (A2M) (Fig. S2), was significantly reduced. However, when using UC only to isolate exosomes from plasma (Fig. 2c, first lane), the highly abundant soluble plasma proteins IgG and albumin remained as contaminants, even though they were less prominent than non-enriched samples or enriched post-exosome fractions (P1-P10). The presence of contaminating abundant plasma proteins in exosome preparations is problematic for downstream proteomics analysis, biasing against detection of low-abundant exosome proteins. Elimination of contaminating proteins is a crucial benchmark for further optimization of exosome isolation.

**Fig. 2.**
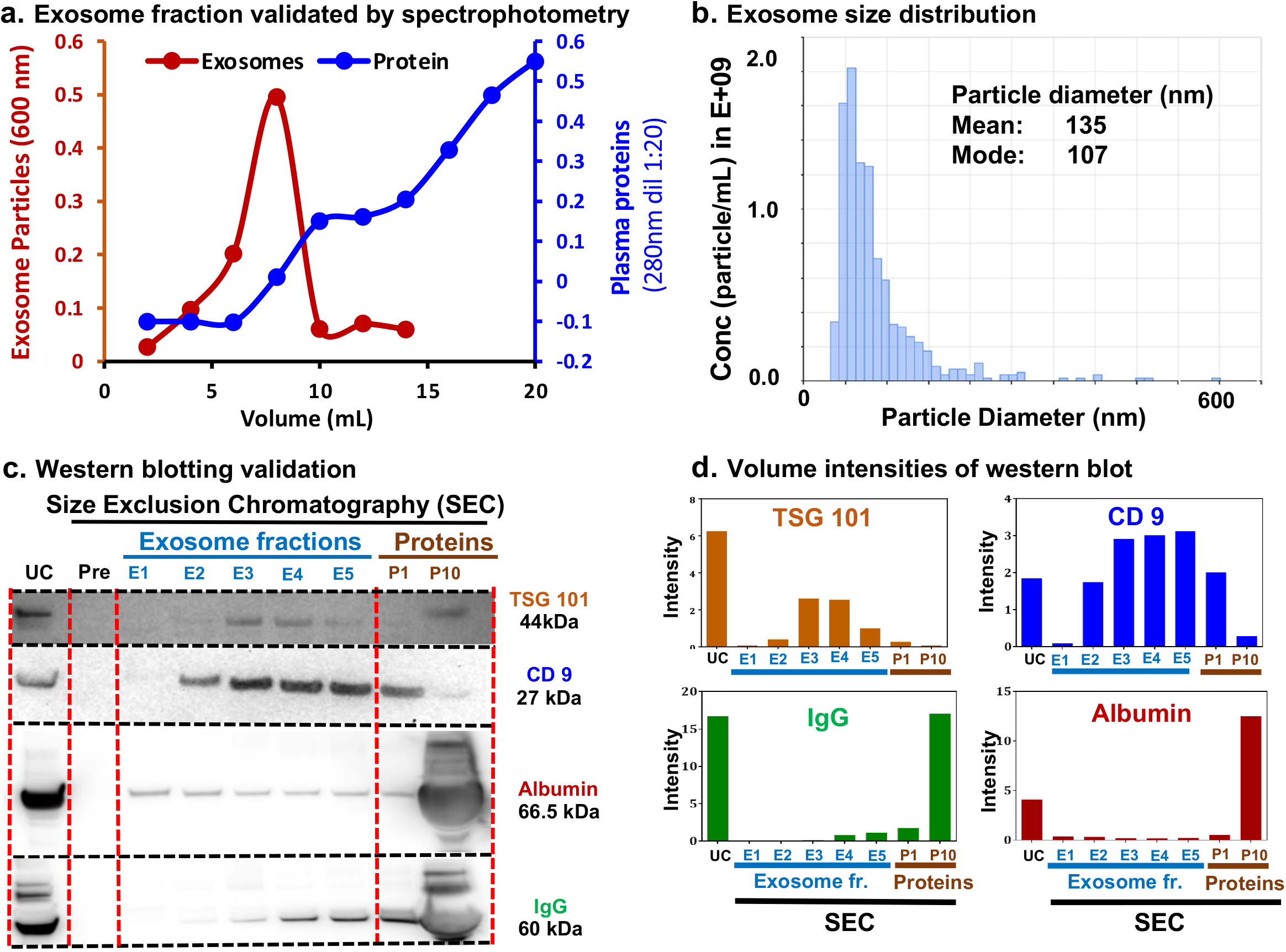
Characterization of SEC-extracted plasma exosomes. (A) Exosome fractions were validated using a spectrophotometer. Red: exosome (OD600), blue: plasma proteins (OD280, dil 1:20). (B) Bar plot showing the exosome size distribution (qNANO GOLD measurements). (C) Western blots confirmed the presence of CD9 and TSG101 proteins (exosome markers) in exosome fractions. Plasma protein contaminants were determined by the presence of IgG and albumin. E1–E5: exosome fraction, P1: pool of fractions 11 + 12, and P10: pool of fractions 29 + 30, 1:20 dilution) were loaded to compare the plasma proteins with the exosome fractions. (D) Semi-quantitative estimation of the volume intensities of western blot bands using Image J software.

Next, we explored SEC fractionation; however, SEC alone leads to relatively high sample volumes (20 mL; 10 mL exosome-containing fractions; E1-E5 + 10 mL protein fractions; P1-P5, Fig. S3). Twenty SEC-extracted plasma fractions (1 mL each) containing intact exosomes were initially analyzed by spectrophotometry as a crude assessment of exosomes. Most plasma exosomes eluted within the first 10 fractions, but smaller plasma proteins eluted more slowly, with maximum concentration in the 20^th^ fraction (Fig. 2a). We pooled two consecutive fractions into one, resulting in a total of five exosomes-enriched fractions (E1–E5), one post-EV fraction (P1), and a pooled soluble protein fraction (P10) containing all subsequent fractions. The details of elution and pooling strategies are provided in Fig. S3. Subsequently, we performed western blotting of the plasma exosome protein fractions (E1–E5) using the exosome specific markers CD9 and TSG101. Fractions E2– E5 showed CD9 (27 kDa) and TSG101 (44 kDa) proteins (Fig. 2c). To rule out contamination by plasma proteins, we tested for two abundant plasma proteins, albumin, and IgG. Soluble plasma protein band intensities in SEC-extracted exosome fractions E1–E5 were drastically lower than those in UC purifications (Fig. 2c), suggesting the high purity of SEC exosome fractions (Fig. 2c).

We further tested combinations of SEC/UC or SEC/UF to concentrate the exosomes and/or further reduce plasma protein contaminations. Using either approach, we obtained even higher purity of plasma exosomes, as determined by particle counting, particle diameter, and the presence of exosome-specific surface protein markers. We estimated the exosome counts to be around 1×10e^10^ exosome particles per mL of plasma by qNano Gold Nanoparticle Characterization instrument (Fig. 2b). Proteomics analysis of the exosomes isolated by SEC/UC or SEC/UF showed similar proteins and exosome numbers (Fig. S4). Thus, SEC coupled to either enrichment strategy results in exosome samples with significantly greater purity than UC alone and without sacrificing yield. SEC/UF was determined to be the most time- and cost-effective workflow, together with good results from initial MS experiments, and was thus used for all further proteomics and miRNA experiments.

### Plasma exosome enrichment for spectral library generation

For detailed quantitative proteomic analysis of exosomes, we developed in-house spectral libraries for plasma samples from five young (20–26 years) and five old (60–66) individuals, using two approaches (Fig. 1). The first approach builds a comprehensive spectral library that serves as a resource for future work, and that can be applied to DIA analysis of human exosome samples, particularly from plasma samples. The second approach was intended to show a robust alternate method that does not require the up-front generation of a deep library by fractionation, which can be time-consuming. A DDA spectral library was generated from DDA acquisitions of the 25 HPRP fractions of pooled plasma exosome proteins, and 2323 protein groups were identified upon processing with Spectronaut (Fig. 3a; Table S2a); among which 1859 proteins were identified with at least two peptides (Table S2b). Next, 10 DIA acquisitions, corresponding to plasma exosome samples from 10 individuals in our cohort, were processed using the directDIA algorithm in Spectronaut generating the DIA-only library consisting of 856 protein groups (Fig. 3a; Table S3a); among which 717 proteins were identified with at least two peptides (Table S3b). The DDA and directDIA libraries showed that 11,436 peptides (Fig. 3b) and 648 proteins (Fig. 3c) overlapped.

**Fig. 3.**
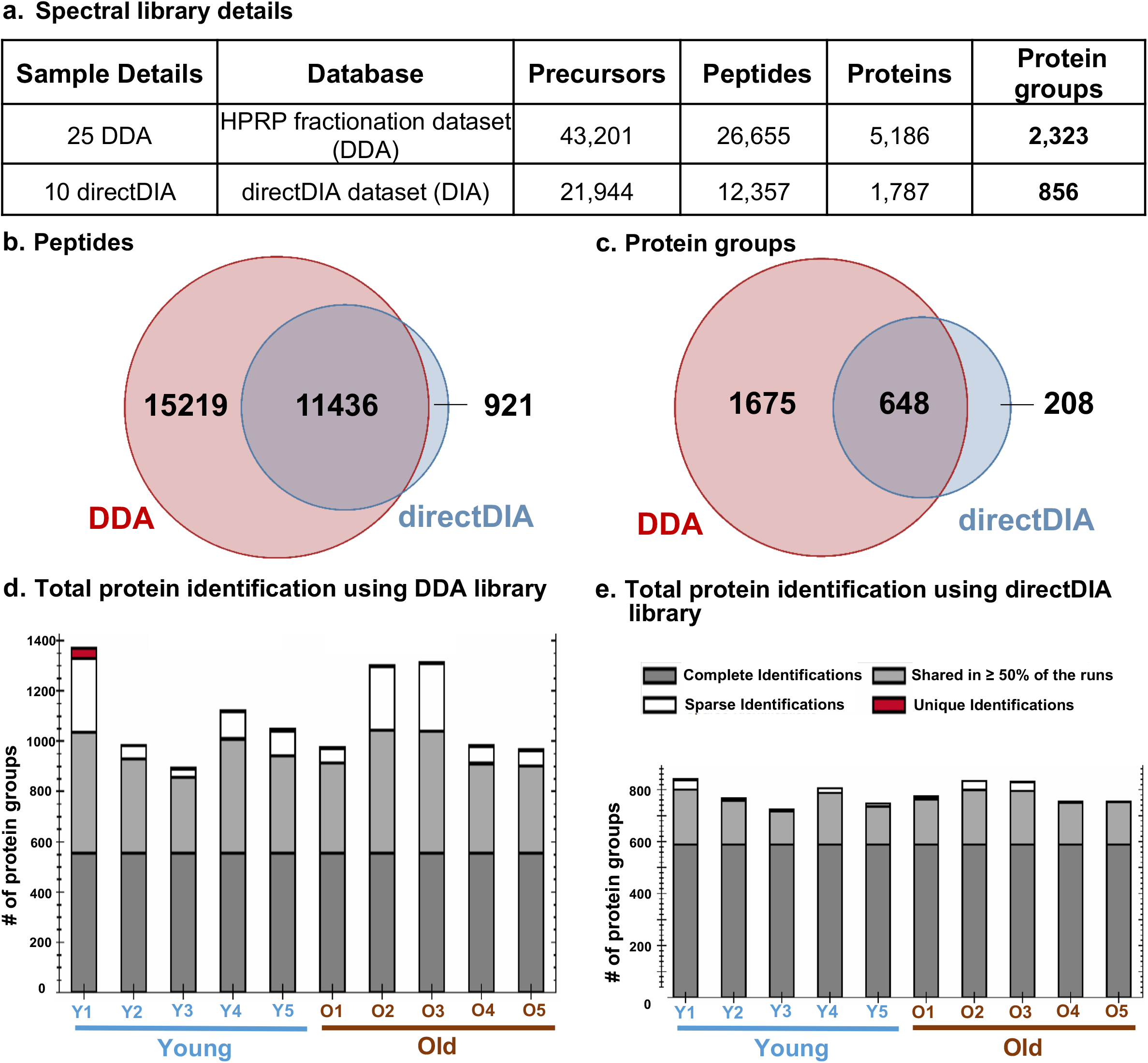
HPRP-fractionation and analysis of plasma exosomes for the generation of DDA and directDIA-MS deep spectral libraries. (A) Table showing the number of precursor ions, peptides, proteins, and unique protein groups in the deep DDA spectral library and in the directDIA spectral library. (B) Venn diagram displaying common and unique peptides in DDA and directDIA library. (C) Venn diagram showing common and unique protein groups in the DDA and directDIA libraries. (D) Bar graphs showing the distribution of individual plasma exosome proteins identified and quantified using either (D) the deep DDA spectral library or (E) the directDIA library.

Given that the fractionated deep-proteome library was most comprehensive and comprised a larger number of proteins, we performed all subsequent bioinformatic assessments using the DDA library. We compared our DDA library with exosome proteins cataloged in the Exocarta database.^50^ About 80% of the exosome DDA spectral library (1485 proteins) overlapped with exosome proteins listed by Plasma Exocarta, confirming a high enrichment of exosome proteins. Interestingly, 374 plasma exosome proteins were newly identified in our study (Fig. S5), potentially representing novel plasma exosome proteins. Interestingly, we found 410 (16%) plasma exosome proteins overlapped with known urinary exosome proteins,^44^ whereas 1,449 (60%) and 611 (24%) proteins were unique to plasma and urine exosomes, respectively (proteins with ≥ 2 unique peptides) (Fig. S6; Table S4).

### Plasma exosome proteins from young and old donor groups

We performed a pilot study using plasma samples from five young donors (20–26 yrs) and five older donors (60–66 yrs) (Table S1) to determine exosome-specific aging signatures. We used DIA-MS protein quantification to analyze young and old plasma exosomes and matched them independently against DDA and directDIA spectral libraries described in method section (Table S5a, Table 6a, Figs. 3d and 3e). We reproducibly identified and quantified a total of 1,318 (Fig. 3d, Table S5b) and 741 (Fig. 3e, Table S6b) proteins with 1% false discovery rate (FDR) and two unique peptides searching against the deep spectral DDA and DIA libraries, respectively. Interestingly, using both the DDA library for DIA quantification as well as in the directDIA workflow, we identified about 600 proteins with complete detection profiles across all individuals. Out of 1,318 exosome proteins identified using the DDA deep spectral library for DIA processing, 804 (60%) were annotated as exosome-related proteins (Table S7). This rather high percentage of exosome-specific proteins in this unbiased proteomics study confirmed the high yield and purity of the extracted plasma exosomes. Partial least squares-discriminant (PLS-DA) analysis clustered young and old individuals into separate groups (Fig. 4a). Volcano plots of Q-value versus log2 ratio of old versus young (old/young) individuals showed significantly altered expression (Q-value <0.05, log-fold change >absolute 0.58) of 144 exosome proteins, of which 92 were down-regulated, whereas 52 were up-regulated in the old cohorts (Fig. 4b; Table S5c). Further, we found 175 significantly altered proteins (Table S6c) when using the directDIA workflow, and 123 significantly changed proteins were common using the DDA and directDIA libraries (Fig. 4c).

**Fig. 4.**
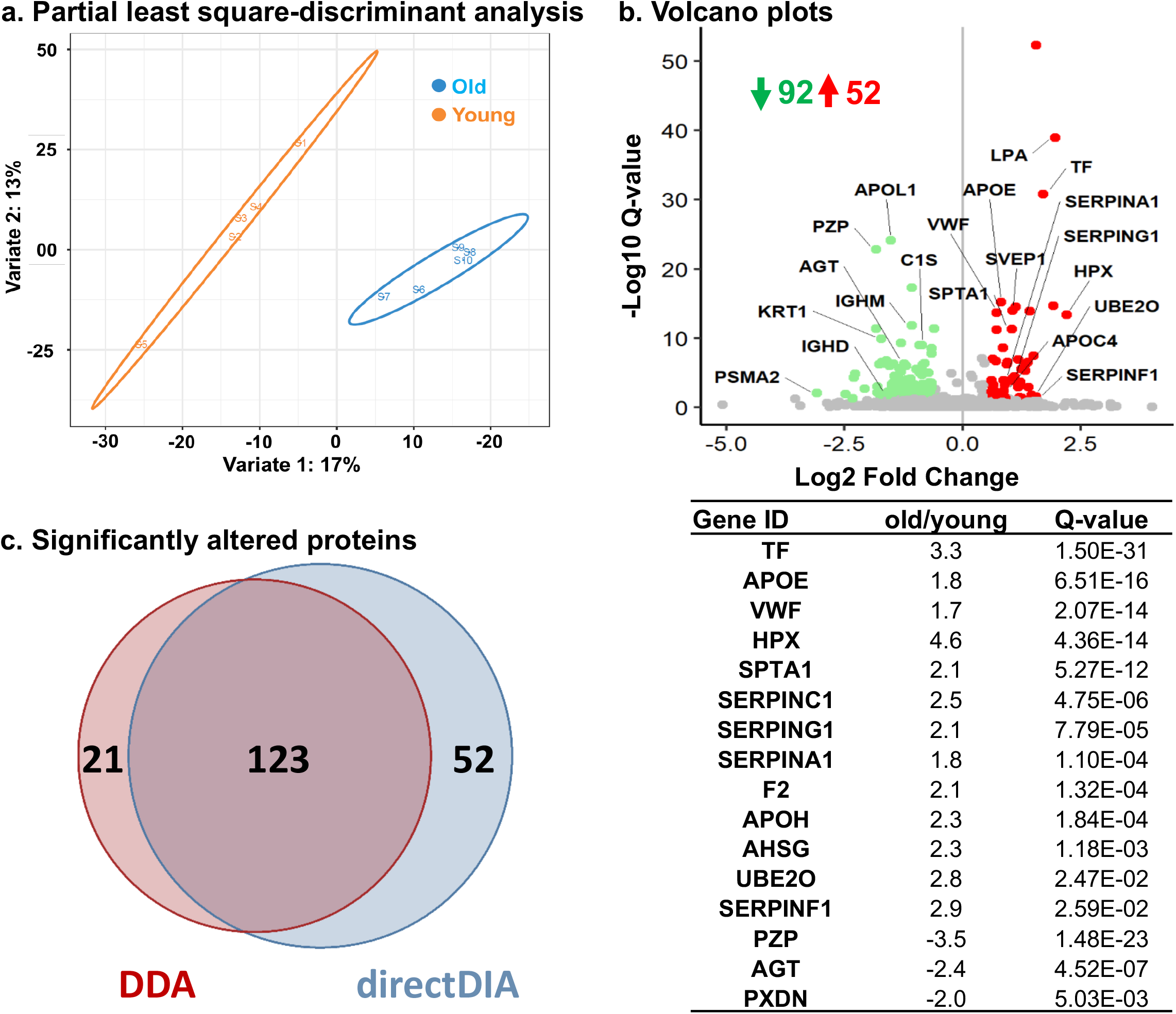
Supervised machine learning classified young (n=5) and aging (n=5) donors based on DIA-MS analysis against DDA deep spectral library in a pilot study. (A) Partial least square-discriminant analysis (PLS-DA) clusters young and aging donor cohorts into two separate groups (B) Volcano plot showing Q-values (−log10) versus protein ratio of (log2 aging vs. young). Green: down-regulated (fold-change < -1.5, Q-value< 0.05), red: up-regulated (fold-change > 1.5, Q-value < 0.05), grey: no significant change. A few selected differentially abundant proteins are labeled. (C) Venn diagram showing common and significantly altered proteins that resulted from the DIA Quantification using directDIA and DDA-MS spectral libraries, respectively. Few of the significantly altered proteins are shown in the Table.

Our deep DDA spectral library was also compared with (i) previously identified radiation (X-ray) and chemically (doxorubicin) induced-senescence-associated exosome proteins and (ii) previously identified urine exosome proteins.^44^ Our plasma exosome protein library overlapped with 242 known senescence exosome proteins, whereas 1617 and 298 proteins were unique to plasma and senescence exosomes, respectively, for proteins with ≥ 2 unique peptides (Fig. 5a; Table S8). Also, the differentially altered exosome proteins identified with the DDA spectral library in this pilot study were compared with (i) the PlasmaExocarta database (Fig. S7) and (ii) the exosome-SASP protein Atlas (*SASPAtlas*.*com*) published by our group (Fig. 5b, Table S9).^3^ We also identified 21 significantly changing plasma exosome proteins (old/young) that are common to the exosome-SASP), suggesting potential senescence-derived and aging-related biomarkers are relevant in the plasma exosome proteome (Fig. 5b; Table S9). This novel line of work for aging research and aging biomarkers will require more extensive validation in future studies. The significantly altered exosome proteins identified in the study were categorized based on subcellular locations, namely extracellular space (98%), membrane (64%), exosome (62%), plasma membrane (42%), cytosol (42%), and cell surface (8%) (Fig. 5c).

**Fig. 5.**
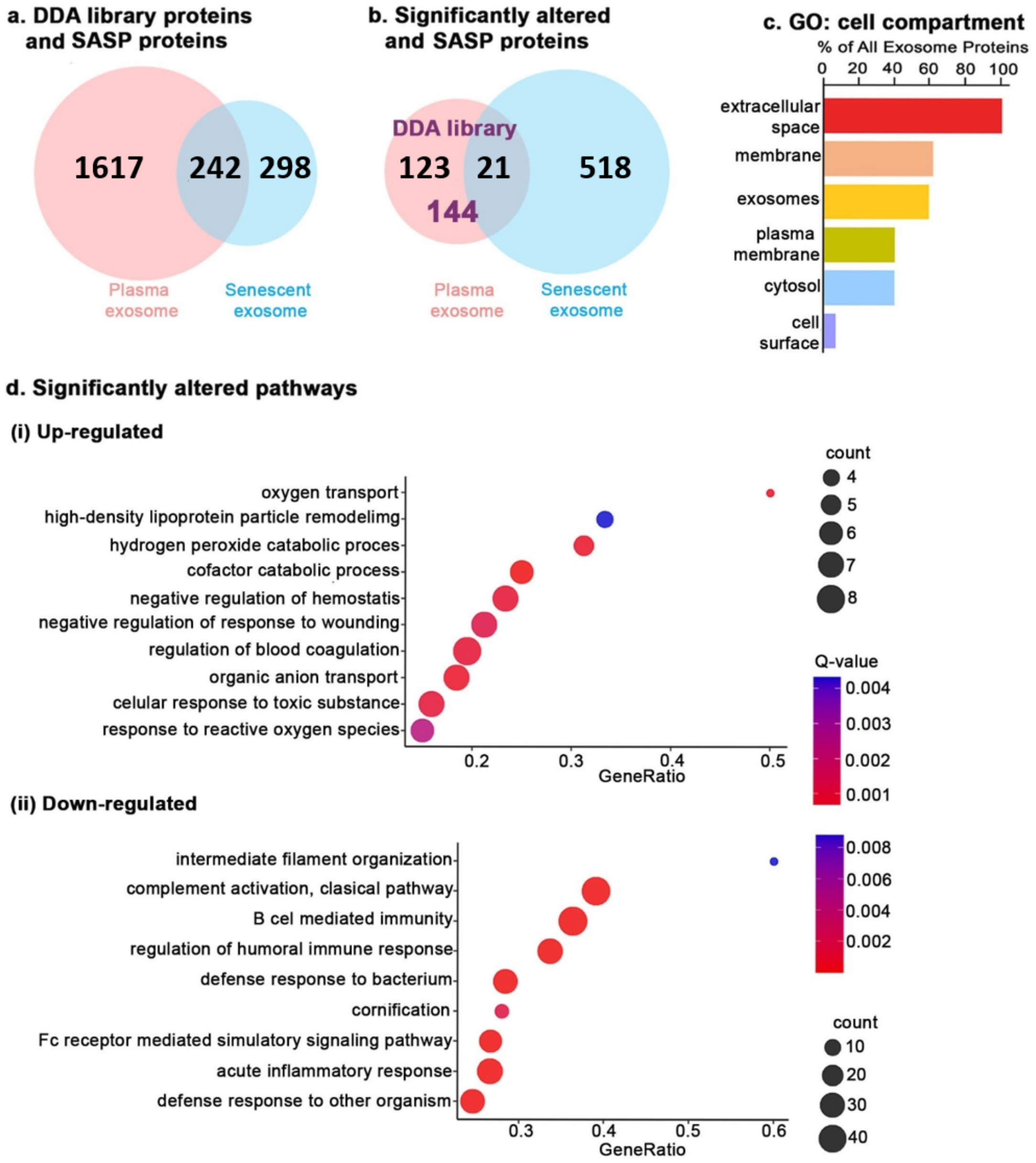
Aging alters exosome features and composition. (A) Venn diagram showing common and unique proteins in DDA library and senescence samples induced by irradiation and chemical treatment.^3^ (B) Venn diagram showing common and unique significantly altered (fold change ≥1.5, Q-value <0.05) proteins in aging and senescence samples. (C) Enrichment analysis of gene-ontology/cellular compartments overrepresented among significantly changing plasma exosome proteins. (D) Pathway and network analysis of significantly regulated plasma exosome proteins in young and aging donors (i) up-regulated and (ii) down-regulated proteins in aging cohorts.

### Age-associated signature exosome proteins and altered pathways

To identify the biological relevance of our results, the differentially expressed exosome proteins from our DIA-MS analysis were integrated and highlighted in all possible ConsensusPathDB-human metabolic pathways using the GO annotation toolbox. Pathway enrichment analysis indicates that plasma lipoprotein particle remodeling, blood coagulation, negative regulation of response to wounding, and reactive oxidant activity were significantly up regulated in plasma exosomes from old donor cohorts. Conversely, defense response to bacteria, acute inflammatory response, humoral immune response regulation, cornification, and intermediate filament cytoskeleton organization was significantly decreased in samples from old donors (Fig. 5d; Table S10).

### Exosome miRNA vary significantly in old and young donor cohorts

To complement the exosome proteomics data, we performed a pilot analysis of plasma exosome miRNA cargo to compare the five samples from young (20–26 yrs) and five from old (60–66 yrs) donors that were also used for the proteomics study (Fig. 1). The miRNA sequences were mostly 18–24 nucleotides (Fig. 6a). In total, 1,375 miRNAs were extracted from our highly pure plasma-derived exosomes (Table S11a), and 331 miRNA were consistent. In fact, 226 out of 331 miRNA were common between exosomes from both young and old individuals, and 88 and 17 were unique to plasma exosomes from old and young donors, respectively (Fig. 6b; Table S11b). Volcano plots of p-value versus log2 fold-change of old vs. young samples showed six miRNA to be significantly altered. PC-3p-73204_81, PC-3p-7719_599, hsa-miR-27a-5p, hsa-miR-874-3p, and PC-3p-8403_561 were up-regulated, and hsa-miR-190a-5p was down-regulated in old donors (Fig. 6c; Table S11c), indicating that these miRNAs could serve as potential biomarkers of aging.

**Fig. 6.**
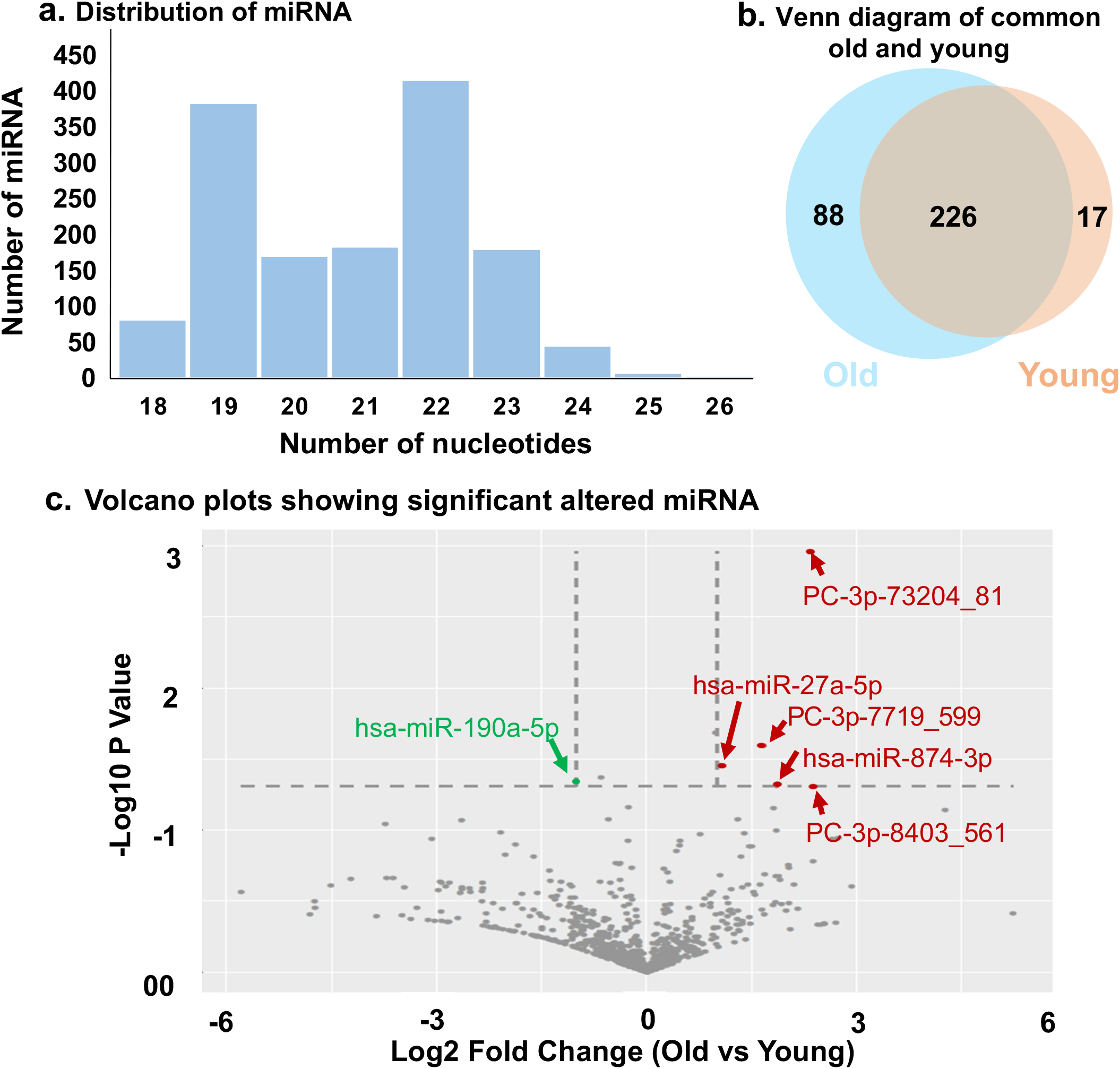
Aging alters plasma exosome miRNA. (A) Bar graph showing the distribution of miRNA. (B) Venn diagram showing common and unique plasma exosome miRNA in young and aging donor cohorts. (C) Volcano plots showing p-values (−log10) versus miRNA ratio of (log2 aging vs. young). Green: down-regulated (fold-change < -1.5, p-value< 0.05), red: up-regulated (fold-change > 1.5, p-value< 0.05), grey: no significant change. Differentially abundant miRNA are annotated.

## Discussion

In the present study, we optimized the isolation and enrichment of plasma exosomes for downstream’ omics’ applications and applied this protocol to dissect the overall changes in plasma exosomes cargo composition during aging. We found that a sequential SEC/UF approach yielded highly pure plasma exosomes, based on exosome particle diameter, the presence of exosome-specific markers, and more importantly, low levels of abundant plasma proteins. We created a deep spectral protein library of plasma exosomes of high purity cataloging ∼2,300 protein that can be applied to plasma biomarker studies in exosomes, particularly to aid in DIA workflows. DIA-MS proteomics analysis enabled quantification of ∼1,300 proteins and suggest over 144 aging biomarker candidates when measuring samples from young and old individuals as part of an aging pilot study. We also identified multiple potential exosome miRNA biomarkers of aging.

Isolation of pure plasma exosomes is challenging but vital for biomarker discovery. Current exosome enrichment approaches rely either on the size of extracellular vesicles or target-specific surface markers (immunoaffinity-based strategies). Here we used a combination of SEC/UF or SEC/UC to obtain highly pure plasma exosomes based on their size. SEC/UC was not used for further proteomics analysis as exosomes are often by the high centrifugal force. We also showed that the optimized method, SEC/UF provides the highest exosome yield and is time-efficient while minimizing contamination from abundant plasma proteins and better than other exosome extraction methods, including UC (Fig. 2).

We generated two high-quality plasma exosome spectral libraries (Fig. 3), based on DDA and directDIA workflows. The comprehensive DDA spectral library for human plasma exosomes will serve as a resource to researchers and clinicians. However, any DDA library preparation can be very time- and resource-consuming, particularly for small projects. We also assessed our workflow for directDIA spectral library. Although we observed fewer proteins in the directDIA spectral library than the DDA library, surprisingly, we identified a similar number of differentially and significantly changing exosome proteins in our pilot study (Fig. 4c). The directDIA approach is particularly useful in cases where no pre-existing large libraries are available, such as in non-model organisms or tissues. The directDIA approach is a highly robust alternative workflow and suitable for analyzing the clinical samples for biomarker discovery.

Biomarker research has expanded from identifying and quantifying proteins for disease conditions to empirical studies of human populations to understanding physiological processes that change with aging and age-related diseases. Senescent cells are one of the hallmarks of aging that drive aging processes via the secretion of various soluble and extracellular vesicle-enclosed proteins collectively known as SASP. ^51, 52^ Our laboratory recently reported an expansive list SASP factors^3, 10^ and extracellular vesicles, including exosomes,^3^ that could serve as biomarkers of aging in human plasma. Borghesan et al. described the role of exosomes in mediating paracrine senescence to the brain and other organs.^53^ Therefore, plasma exosomes are potential drivers and biomarkers of aging. We identified 144 proteins significantly associated with aging in plasma-derived exosomes from humans, including proteins positively and negatively correlated with aging. Interestingly, numerous variants of immunoglobulins (n=52, including heavy and light chains, as presented in Table S5c) showed a strong negative correlation with aging. The up-regulation of hemopexin (HPX) in the exosomes of old donor cohorts is consistent with the accumulation of HPX in CSF samples from Alzheimer’s disease (AD) patients. ^54^ Several significantly altered proteins (n=21) associated with well-known aging processes or the SASP, such as serine protease inhibitors (SERPIN family) SERPINA1, SERPINC1, pregnancy zone protein (PZP), prothrombin (F2), and peroxidasin (PXDN) overlapped with old plasma exosome proteins (Table S10). Notably, various SERPINs variants appear to be prominent plasma biomarkers of aging^55^ and the SASP.^3^ We also found that several components of these pathways, including acute inflammatory response (alpha-2-HS-glycoprotein (AHSG), F2, HP, SERPINA1, SERPINC1), humoral immune response (F2, HPX, IGHG2, IGHG4, SERPING1),^56^ and coagulation, were indeed up-regulated in the samples from old donors. These pathways have well-documented links to aging^56^ or senescence.^3, 10, 55, 57^

Moreover, the up-regulation of beta-2-glycoprotein 1 (APOH), F2, KNG1, SERPINC1, SERPING1, von Willebrand factor (VWF), serotransferrin (TF), and proteins (related to complement and coagulation pathways) in samples from old donors (Fig. 4, Table S10) agrees with a report that suggests an increased risk for thrombosis as a common age-associated complication.^52^ Our group previously showed that activated complement and coagulation pathways due to the SASP increased the risk of age-related complications.^10^ Also, previous studies have reported that proteins can be increasingly ubiquitinated during aging and in age-related diseases, such as AD.^58, 59^ In this study, we detected an up-regulation of E2 ubiquitin-conjugating enzyme (UBE2O), a component of the ubiquitin system, in samples from old donors. Down-regulation of proteasome subunit alpha 2 (PSMA2), which are proteasome subunits, could be associated with the accumulation of damaged and misfolded proteins, yet another hallmark of aging.^60^ Declined proteasome activity is linked to age-related neurodegenerative disorders, such as AD, Parkinson’s disease, and Huntington’s disease.^61^ Finally, the down-regulation of different variants of heavy and light chain immunoglobulins, which are antibody components, suggests reduced immunogenic potentials during aging.

We also investigated miRNAs, which are well-known components of exosome cargo. These short non-coding RNAs (approximately 18–25 nucleotides in length) bind mRNAs through partial base-pair complementarity with their target genes, resulting in post-transcriptional regulation. Here, we reproducibly identified 331 miRNAs from pure plasma-derived exosomes. Of these, 226 miRNA were detected in both young and old donors, and 88 and 17 were unique to plasma exosomes from old and young donors, respectively. Two miRNAs, hsa-miR-199b-5p and hsa-miR-409-3p, out of 88 miRNAs unique to the aging individuals were also reported in a patent granted to Dr. Judith Campisi (WO2019002265A1).^62^ These specific miRNAs could be developed into future senescence biomarkers. miRNAs affect SASP factors and may contribute to profound changes in SASP expression profiles.^63^ Evidence of age-associated changes in miRNA expressions was also seen in various models, ranging from nematodes to humans.^64^ Likewise, in the present study, six miRNA, including miR-27a, miR-190a, and miR-874, were differentially regulated in young and old cohorts. In degenerative diseases such as osteoarthritis^65^ and AD,^66^ there is an up-regulation of miR-27a, while its dysregulation was reported in different cancer types.^67^ Also, miR-190 dysregulation was detected during cancer development and progression^68^ and in plasma neural-derived exosomes of AD.^66^ Finally, miR-874 was identified in circulating brain fluid from patients with mild cognitive impairment^69^ and as a biomarker for the early stage of AD development.^70, 71^ Thus, the role of miR-27a, miR-190, and miR-874 in age-related diseases and their dysregulation in our study cohorts strongly suggest their potentially important role during aging. Further investigations with larger donor groups and longitudinal analysis are needed to confirm the potential of these specific miRNAs as diagnostic and/or prognostic biomarkers.

## Experimental

### Materials

HPLC-grade acetonitrile and water were obtained from Burdick & Jackson (Muskegon, MI). Reagents for protein chemistry including iodoacetamide, dithiothreitol (DTT), ammonium bicarbonate, formic acid (FA), and urea were purchased from Sigma Aldrich (St. Louis, MO). Sequencing grade trypsin was purchased from Promega (Madison, WI). HLB Oasis SPE cartridges were purchased from Waters (Milford, MA).

### Procurement of plasma samples and storage

Plasma samples (n=10) of two different age groups (young: 20–26 yrs; old: 60–66 yrs) were obtained from the Blood Centers of the Pacific (San Francisco, CA) as detailed in Table S1 (no IRB approvals were necessary for these sample sets). Plasma samples were aliquoted and stored at -80°C until further use.

### Exosome extraction

Plasma exosomes were pre-processed and isolated from 2 mL of human plasma by size exclusion chromatography (SEC), using qEV columns (#qEV35, IZon), according to the manufacturer’s directions. Briefly, the following steps were performed: 2 mL of human plasma was centrifuged at 3,000 x g for 10 min to remove cell debris. The supernatant was again centrifuged at 12,000 x g for 30 min to remove apoptotic cell bodies, and the extracellular vesicles remained in the supernatant. The cleared supernatant was passed through the qEV size-exclusion column: the column was maintained at room temperature for 20 min to reach operational temperature before use. The column was activated by flushing 1.5x column volume (∼60 mL) of degassed phosphate-buffered saline (PBS). Plasma supernatant was passed through the activated qEV size exclusion columns (drip by gravity). Different fractions were collected by filling-up the reservoir with PBS buffer (50 mL). Initial 14 mL of flowthrough was the void volume (Fig. S3), the subsequent 10 mL (collected in 5 x 2 mL centrifuge tubes: labelled as E1–5) of flowthrough was considered as an enriched exosome fraction, based on the western blot analysis with the exosome-specific protein markers (Fig. 2). The remaining 26 mL were marked as the post-exosome fractions that contained plasma proteins (Fig. S3). qEV size-exclusion exosome fraction (E1–5) was quality-checked spectrophotometrically (#SpectraMax Plus 384, Molecular devices), where OD_600_ was used for exosome quantification and OD_280_ for plasma proteins. Based on higher plasma protein contamination in fraction E5, we pooled only fractions E1–E4 to obtain highly pure exosomes. Subsequently, fractions E1–E4 were concentrated by ultrafiltration (UF, Amicon® Ultra 50 kDa) or ultracentrifugation (#Optima, The Beckman Coulter Life Sciences) for all further proteomics and miRNA analysis.

### Size distribution analysis

Particle diameter was assessed by tunable resistive pulse sensing (TRPS) on an IZON qNano Nanoparticle Characterization instrument using an NP150 nanopore membrane at calibration with 110-nm carboxylated polystyrene beads at a concentration of 1 × 10^10^ particles/mL (Zen-bio, Research Triangle, NC).

### Western blotting

Aliquots of 20⍰μg of protein/well were separated on 12% SDS-PAGE and transferred onto PVDF membranes under semi-dry conditions using the Trans-Blot Turbo Transfer System transfer unit (Bio-Rad, 1704150). After transfer, equal loading of the samples in each lane was confirmed by Ponceau S staining (Sigma-Aldrich, P7170-1L). Western blot analysis was performed using 1:1000 dilution of CD9 (Abcam, Ab96696), TSG101 (Santa Cruz Biotechnology, sc-7964), albumin (Abcam, Ab106582), or IgG (Abcam, Ab200699), and anti-rabbit HRP-conjugated antibody (Bio-Rad, 1706515) or anti-mouse HRP-conjugated antibody (Abcam, Ab6728) as a secondary antibody (1:3000 dilution). Blots were developed as per the manufacturer’s protocol with the Pierce ECL Western Blotting Substrate (ThermoFisher Scientific, 32106). Imaging was performed by Azure Biosystems c600. Densitometric analysis of bands was performed using Image J software (IJ 1.46r; NIH), and intensities obtained were expressed as mean⍰±⍰standard deviation.

### Sample processing for MS analysis

#### Exosome protein extraction and trypsin digestion

Plasma exosome samples were lysed in lysis buffer (8 M urea, 0.1 M ammonium bicarbonate), and aliquots were reduced by treatment with 10 mM tris(2-carboxyethyl) phosphine (TCEP) and 40 mM 2-chloroacetamide (CAA) for 1 h at 37°C. The solution was diluted with 0.1 M ammonium bicarbonate buffer to lower the urea concentration to 1.5 M, and digested with trypsin (Promega, Madison, WI) at 1:100 ratio overnight at 37°C. Peptides were purified using MacroSpin clean-up columns (NEST group, Southborough, MA), following the manufacturer’s protocol. Eluted peptides were vacuum dried until complete dryness and resuspended in 60 µl buffer A (1% acetonitrile acidified with 0.1% formic acid) spiked with iRT peptides (Biognosys, Schlieren, Switzerland). Peptides were quantified using nano-drop (#BMG Labtech, Spectrostar Nano) and adjusted to a 1 mg/mL concentration.

#### High pH reversed-phase fractionation for deep spectral library generation

For DDA library generation, 400 µg of exosome peptides (pooled from 10 different plasma exosome samples) were injected onto Dionex Ultimate 3000 LC (Thermo Scientific) coupled to ACQUITY UPLC CSH1.7 mm C18 column (2.1 x 150 mm) (Waters, Milford) for fractionation using high pH reversed-phase (HPRP) fractionation. Peptides were chromatographically separated in a 30-min non-linear gradient from 1% to 40% HPRP buffer B (Buffer B: 100% ACN; Buffer A: 20 mM ammonium formate, pH 10). A micro fraction was taken every 45 seconds and pooled into 24 final peptide fractions, vacuum dried, resuspended in 20 mL buffer A spiked with indexed retention time (iRT peptides), and quantified. Each fraction was acquired in DDA mode as described below.

### Mass spectrometric data acquisition

2 µg of digested exosome samples were separated using liquid chromatography coupled to a mass spectrometer. The analytical column was in-house packed into a fritted tip emitter to a length of 50 cm (ID 75 µm) (New Objective, Woburn, MA) using the CSH C18 phase (1.7 µm) (Waters, Milford, MA). The column was operated using an Easy nLC 1200 (Thermo Fisher Scientific, San Jose, CA) coupled online to an Exploris 480 spectrometer (Thermo Fisher Scientific). Peptides were eluted at 250 ml min^-1^ using a non-linear 2-h gradient from 1% to 45% buffer B (Buffer B: 80% ACN+ 0.1% FA; Buffer A: 0.1% FA).

For DDA library generation based on the HPRP samples, the mass spectrometer was operated in a data-dependent mode. Briefly, the following settings were used: MS1 scan resolution: 60,000; MS1 AGC target: 300; MS1 maximum IT: 25 ms; MS1 scan range: 350–1650 Th; MS2 scan resolution: 15,000; MS2 AGC target: 200; MS2 maximum IT: 25 ms; isolation window: 4 Th; first fixed mass: 200 Th; NCE: 27; minimum AGC target: 1e3; only charge states 2 to 6 considered; peptide match: preferred; dynamic exclusion time: 30 s.

In DIA mode all samples were acquired in a randomized fashion with regards to the different conditions (young, old). For DIA, the mass spectrometer was operated using the following parameters for the MS1 scan: scan range: 350 to 1650 Th; AGC target: 300%; max injection time: 20 ms; scan resolution: 120,000. The MS1 was followed by targeted MS2 scan events with the following settings: AGC target: 1000%; max injection time: 54 ms; scan resolution: 30,000; scan range: 250-1000 Th; normalized collision energy: 27. The number of DIA segments and the segment widths were adjusted to the precursor density and to achieve optimal data points across each peak for each acquisition (Supplementary Table 12). The DIA methods consisted of 40 DIA segments and had a cycle time of 3.2 s.

### Spectral library generation from DDA and DIA

Two spectral libraries were generated i) a deep DDA-based spectral library, and ii) a DIA-only spectral library (directDIA) (Fig. 1). The DDA deep spectral library was generated by performing DDA analysis on 25 HPRP fractions, representing fractions from a pooled mixture of peptides obtained from the same 10 plasma exosome protein samples (from five young and five old individuals). DDA raw files (n = 25) were individually searched with SpectroMine 2.0.200228.46524 (Biognosys) against the Human UniProt FASTA including isoforms (downloaded on January 1, 2020, containing 68674 proteins) using the following settings: fixed modification: carbamidomethyl (C); variable modifications: acetyl (protein N-term), oxidation (M); enzyme: trypsin/P with up to two missed cleavages. Mass tolerances were automatically determined by SpectroMine, and other settings were set to default. Search results were filtered using a 1% false discovery rate on the precursor ion, peptide, and protein levels. The plasma exosome spectral libraries were generated using the default values in SpectroMine.^72, 73^ To generate directDIA spectral libraries, 10 individual DIA acquisitions (from five young and five old individuals) were used to generate spectral libraries using the directDIA algorithm in Spectronaut version: 12.0.20491.18.30559 directly searching the DIA data against a human FASTA file. All Spectronaut details and parameter settings can be found as uploads in the data repository (see data accession below).

### Quantification and DIA processing

Quantitative and statistical analysis was performed processing protein peak areas determined by the Spectronaut software. Prior to library-based analysis of the DIA data, the DIA raw files were converted into htrms files using the htrms converter (Biognosys). MS1 and MS2 data were centroided during conversion, and the other parameters were set to default. The htrms files were analyzed with Spectronaut (version: 14.0.200601.47784, Biognosys) using the previously generated libraries and default settings to perform quantitative data analysis with the two libraries generated (see above): the directDIA library and the deep DDA spectral library. Briefly, Calibration was set to non-linear iRT calibration with precision iRT selected. DIA data was matched against the above-described spectral library supplemented with decoys (library size fraction of 0.1), using dynamic mass tolerances. The DIA data was processed for relative quantification comparing peptide peak areas from different conditions (old versus young). For the DIA MS2 data sets, quantification was based on XICs of 3-6 MS/MS fragment ions, typically y- and b-ions, matching to specific peptides present in the spectral library. Interference correction was enabled on MS1 and MS2 levels. Precursor and protein identifications were filtered to 1% FDR, estimated using the mProphet algorithm,^74^ and iRT profiling was enabled. Quantification was normalized to local total ion chromatogram. Statistical comparison of relative protein changes was performed with paired t-tests. Finally, proteins identified with less than two unique peptides were excluded from the assay. The significance level was q-value less than 0.05%, and log2 ratio more than 0.58. Both DIA analysis results were filtered within Spectronaut by a 1% false discovery rate on peptide and protein level using a target-decoy approach, which corresponds to a Q value <0.01.

### MS raw data and plasma exosome library availability

All mass spectrometry raw files, spectral libraries, Pilot study results table (Supplementary Table S5), DIA results table (Supplementary Table S6), Spectronaut files, descriptive methods, and other supplementary tables and data have been deposited as MassIVE ID MSV000086782 to the MassIVE repository at ftp://MSV000086782@massive.ucsd.edu (user: MSV000086782, password: winter, preferred engine Firefox) and to the ProteomeXchange Consortium under PXD023897 (http://proteomecentral.proteomexchange.org). The Spectronaut projects that are uploaded to the repositories can be viewed using the free Spectronaut viewer (www.biognosys.com/technology/spectronaut-viewer).

### Pathway and network analysis

Gene set over-representation analyses were performed using the ConsensusPathDB-human tool, release 34 (15.01.2019) on a list of all significantly (q-value < 0.05) increased or decreased proteins in old and young plasma exosomes. Curated pathways for enrichment analysis were referenced from the Gene Ontology (GO) Database with “Biological Process” term categories. Pathways with a minimum of three observed proteins, Q-value < 0.01, and GO terms restricted to levels 4–5 were considered significant. A list of all proteins present in the deep fractionated DDA spectral library was used for background reference. Top selected pathways are plotted in Fig. 5d. Tables containing the entire list of differentially regulated pathways, observed corresponding proteins, reference pathway annotations, statistics, and source databases are included as a supplemental file (Table S10). Dotplot representations of pathway analysis were generated in R with the “ggplot2” package.^75^

### Plasma exosome miRNA extraction and analysis

For plasma exosome miRNA analysis, the extracted plasma exosome samples were outsourced to LC Sciences, LLC (Houston, TX). Briefly, miRNA was extracted from plasma exosomes using the following kits (Cat# RS-930, Illumina). A small RNA (sRNA) library was generated for 10 plasma exosome samples (five young and five old individuals) using the Illumina Truseq™ Small RNA Preparation kit, according to Illumina’s TruSeq™ Small RNA Sample Preparation Guide. The purified cDNA library was used for cluster generation on Illumina’s Cluster Station and then sequenced on Illumina GAIIx following the vendor’s instruction for operating the instrument. Raw sequencing reads (40 nts) were obtained using Illumina’s Sequencing Control Studio software version 2.8 (SCS v2.8), following real-time sequencing image analysis and base-calling by Illumina’s Real-Time Analysis version 1.8.70 (RTA v1.8.70). The extracted sequencing reads were used in the standard data analysis. A proprietary pipeline script, ACGT101-miR v4.2 (LC Sciences), was used for sequencing data analysis.^76^

## Conclusions

Our data suggest an optimized workflow for high purity exosome isolation from human plasma consisting of SEC/UF for downstream biomarker studies. To further facilitate biomarker discovery from plasma exosomes, we created a deep spectral library as a resource for DIA-MS studies. We also showed that this workflow can be useful for identifying protein and miRNA biomarker candidates in a pilot DIA-MS proteomics and miRNA analysis of the exosomes from young and old donors. Our study suggests that exosome proteins could serve as potential aging biomarkers.

## Author Contributions

Conceptualization, BS, JC; experiments and methodology, SKP, RB, LR; data processing, SKP, RB, LR, NB, JB; writing—original draft, SKP, NB, PYD, RB; Figures, SKP, JB, BS; writing—review and editing, BS, JC, RB, LR.

## Acknowledgements

This work was supported by grants from the National Institutes of Health under award numbers U01 AG060906 and U01 AG060906-02S1 (BS), P01 AG017242 and R01 AG051729 (JC), and K99 AG065484 (NB). SKP is supported by a Glenn Fellowship in Aging Research from the Glenn Foundation. We thank John Carroll for graphical support and LC Sciences for miRNA Sequencing Services.

## References

1. J. O. Brett and T. A. Rando, Current Opinion in Genetics & Development, 2014, 26, 33–40.

2. L. Ferrucci, F. Giallauria and J. M. Guralnik, Radiologic Clinics of North America, 2008, 46, 643–652.

3. N. Basisty, A. Kale, O. H. Jeon, C. Kuehnemann, T. Payne, C. Rao, A. Holtz, S. Shah, V. Sharma, L. Ferrucci, J. Campisi and B. Schilling, PLOS Biology, 2020, 18, e3000599.

4. J. Campisi and L. Robert, Interdiscip Top Gerontol, 2014, 39, 45–61.

5. M. J. Yousefzadeh, J. Zhao, C. Bukata, E. A. Wade, S. J. McGowan, L. A. Angelini, M. P. Bank, A. U. Gurkar, C. A. McGuckian, M. F. Calubag, J. I. Kato, C. E. Burd, P. D. Robbins and L. J. Niedernhofer, Aging Cell, 2020, 19, e13094.

6. B. G. Childs, M. Durik, D. J. Baker and J. M. van Deursen, Nature Medicine, 2015, 21, 1424–1435.

7. J.-P. Coppé, C. K. Patil, F. Rodier, Y. Sun, D. P. Muñoz, J. Goldstein, P. S. Nelson, P.-Y. Desprez and J. Campisi, PLoS Biology, 2008, 6, 2853–2868.

8. N. Basisty, A. Kale, S. Patel, J. Campisi and B. Schilling, Expert Review of Proteomics, 2020, 17, 297–308.

9. I. M. Conboy, M. J. Conboy and J. Rebo, Aging, 2015, 7, 754–765.

10. C. D. Wiley, S. Liu, C. Limbad, A. M. Zawadzka, J. Beck, M. Demaria, R. Artwood, F. Alimirah, J.-A. Lopez-Dominguez, C. Kuehnemann, S. R. Danielson, N. Basisty, H. G. Kasler, T. R. Oron, P.-Y. Desprez, S. D. Mooney, B. W. Gibson, B. Schilling, J. Campisi and P. Kapahi, Cell Reports, 2019, 28, 3329–3337 e3325.

11. J. A. Fafián-Labora and A. O’Loghlen, Trends in Cell Biology, 2020, 30, 628–639.

12. O. H. Jeon, C. Kim, R.-M. Laberge, M. Demaria, S. Rathod, A. P. Vasserot, J. W. Chung, D. H. Kim, Y. Poon, N. David, D. J. Baker, J. M. van Deursen, J. Campisi and J. H. Elisseeff, Nature Medicine, 2017, 23, 775–781.

13. M. D’Anca, C. Fenoglio, M. Serpente, B. Arosio, M. Cesari, E. A. Scarpini and D. Galimberti, Frontiers in Aging Neuroscience, 2019, 11, 232.

14. D.-S. Choi, D.-K. Kim, Y.-K. Kim and Y. S. Gho, Proteomics, 2013, 13, 1554–1571.

15. L. Barile and G. Vassalli, Pharmacology & Therapeutics, 2017, 174, 63–78.

16. T. Huang and C.-X. Deng, International Journal of Biological Sciences, 12019, 15, 1–11.

17. M. Rodrigues, J. Fan, C. Lyon, M. Wan and Y. Hu, Theranostics, 2018, 8, 2709–2721.

18. P. A. Gonzales, T. Pisitkun, J. D. Hoffert, D. Tchapyjnikov, R. A. Star, R. Kleta, N. S. Wang and M. A. Knepper, Journal of the American Society of Nephrology, 2009, 20, 363–379.

19. T. Pisitkun, R. F. Shen and M. A. Knepper, Proceedings of the National Academy of Sciences, 2004, 101, 13368–13373.

20. C. Admyre, J. Grunewald, J. Thyberg, S. Gripenbäck, G. Tornling, A. Eklund, A. Scheynius and S. Gabrielsson, European Respiratory Journal, 2003, 22, 578–583.

21. A. Poliakov, M. Spilman, T. Dokland, C. L. Amling and J. A. Mobley, The Prostate, 2009, 69, 159–167.

22. C. Admyre, S. M. Johansson, K. R. Qazi, J.-J. Filén, R. Lahesmaa, M. Norman, E. P. A. Neve, A. Scheynius and S. Gabrielsson, The Journal of Immunology, 2007, 179, 1969–1978.

23. C. Looze, D. Yui, L. Leung, M. Ingham, M. Kaler, X. Yao, W. W. Wu, R.-F. Shen, M. P. Daniels and S. J. Levine, Biochemical and Biophysical Research Communications, 2009, 378, 433–438.

24. S. Keller, C. Rupp, A. Stoeck, S. Runz, M. Fogel, S. Lugert, H. D. Hager, M. S. Abdel-Bakky, P. Gutwein and P. Altevogt, Kidney International, 2007, 72, 1095–1102.

25. M. W. Graner, O. Alzate, A. M. Dechkovskaia, J. D. Keene, J. H. Sampson, D. A. Mitchell and D. D. Bigner, The FASEB Journal, 2008, 23, 1541–1557.

26. M. P. Bard, J. P. Hegmans, A. Hemmes, T. M. Luider, R. Willemsen, L.-A. A. Severijnen, J. P. van Meerbeeck, S. A. Burgers, H. C. Hoogsteden and B. N. Lambrecht, American Journal of Respiratory Cell and Molecular Biology, 2004, 31, 114–121.

27. M. Monguió-Tortajada, M. Morón-Font, A. Gámez-Valero, L. Carreras-Planella, F. E. Borràs and M. Franquesa, Current Protocols in Stem Cell Biology, 2019, 49, e82.

28. H. K. Skalnikova, B. Bohuslavova, K. Turnovcova, J. Juhasova, S. Juhas, M. Rodinova and P. Vodicka, Proteomes, 2019, 7, 17.

29. G. Rabinowits, C. Gerçel-Taylor, J. M. Day, D. D. Taylor and G. H. Kloecker, Clinical Lung Cancer, 2009, 10, 42–46.

30. J. Silva, V. Garcia, A. Zaballos, M. Provencio, L. Lombardia, L. Almonacid, J. M. Garcia, G. Dominguez, C. Pena, R. Diaz, M. Herrera, A. Varela and F. Bonilla, European Respiratory Journal, 2010, 37, 617–623.

31. Y. Tanaka, H. Kamohara, K. Kinoshita, J. Kurashige, T. Ishimoto, M. Iwatsuki, M. Watanabe and H. Baba, Cancer, 2013, 119, 1159–1167.

32. A. Abramowicz, P. Widlak and M. Pietrowska, Molecular BioSystems, 2016, 12, 1407–1419.

33. M. Colombo, G. Raposo and C. Théry, Annual Review of Cell and Developmental Biology, 2014, 30, 255–289.

34. A. Cvjetkovic, J. Lötvall and C. Lässer, Journal of Extracellular Vesicles, 2014, 3.

35. D. W. Greening, R. Xu, H. Ji, B. J. Tauro and R. J. Simpson, Methods Mol Biol, 2015, 1295, 179–209.

36. F. Momen-Heravi, L. Balaj, S. Alian, P.-Y. Mantel, A. E. Halleck, A. J. Trachtenberg, C. E. Soria, S. Oquin, C. M. Bonebreak, E. Saracoglu, J. Skog and W. P. Kuo, Biological Chemistry, 2013, 394, 1253–1262.

37. C. Théry, S. Amigorena, G. Raposo and A. Clayton, Curr Protoc Cell Biol, 2006, Chapter 3, Unit 3 22.

38. I. Helwa, J. Cai, M. D. Drewry, A. Zimmerman, M. B. Dinkins, M. L. Khaled, M. Seremwe, W. M. Dismuke, E. Bieberich, W. D. Stamer, M. W. Hamrick and Y. Liu, Plos One, 2017, 12, e0170628.

39. R. E. Lane, D. Korbie, W. Anderson, R. Vaidyanathan and M. Trau, Scientific Reports, 2015, 5, 7639.

40. R. J. Lobb, M. Becker, S. Wen Wen, C. S. F. Wong, A. P. Wiegmans, A. Leimgruber and A. Möller, J Extracell Vesicles, 2015, 4, 27031.

41. D. D. Taylor, W. Zacharias and C. Gercel-Taylor, Methods Mol Biol, 2011, 728, 235–246.

42. T. Baranyai, K. Herczeg, Z. Onódi, I. Voszka, K. Módos, N. Marton, G. Nagy, I. Mäger, M. J. Wood, S. El Andaloussi, Z. Pálinkás, V. Kumar, P. Nagy, Á. Kittel, E. I. Buzás, P. Ferdinandy and Z. Giricz, Plos One, 2015, 10, e0145686.

43. A. N. Böing, E. van der Pol, A. E. Grootemaat, F. A. W. Coumans, A. Sturk and R. Nieuwland, Journal of Extracellular Vesicles, 2014, 3.

44. S. Guan, H. Yu, G. Yan, M. Gao, W. Sun and X. Zhang, Journal of Proteome Research, 2020, 19, 2217–2225.

45. T. A. Addona, S. E. Abbatiello, B. Schilling, S. J. Skates, D. R. Mani, D. M. Bunk, C. H. Spiegelman, L. J. Zimmerman, A.-J. L. Ham, H. Keshishian, S. C. Hall, S. Allen, R. K. Blackman, C. H. Borchers, C. Buck, H. L. Cardasis, M. P. Cusack, N. G. Dodder, B. W. Gibson, J. M. Held, T. Hiltke, A. Jackson, E. B. Johansen, C. R. Kinsinger, J. Li, M. Mesri, T. A. Neubert, R. K. Niles, T. C. Pulsipher, D. Ransohoff, H. Rodriguez, P. A. Rudnick, D. Smith, D. L. Tabb, T. J. Tegeler, A. M. Variyath, L. J. Vega-Montoto, Å. Wahlander, S. Waldemarson, M. Wang, J. R. Whiteaker, L. Zhao, N. L. Anderson, S. J. Fisher, D. C. Liebler, A. G. Paulovich, F. E. Regnier, P. Tempst and S. A. Carr, Nature Biotechnology, 2009, 27, 633–641.

46. J. D. Chapman, D. R. Goodlett and C. D. Masselon, Mass Spectrometry Reviews, 2013, 33, 452–470.

47. G. Rosenberger, C. C. Koh, T. Guo, H. L. Röst, P. Kouvonen, B. C. Collins, M. Heusel, Y. Liu, E. Caron, A. Vichalkovski, M. Faini, O. T. Schubert, P. Faridi, H. A. Ebhardt, M. Matondo, H. Lam, S. L. Bader, D. S. Campbell, E. W. Deutsch, R. L. Moritz, S. Tate and R. Aebersold, Scientific Data, 2014, 1, 140031.

48. C.-C. Tsou, D. Avtonomov, B. Larsen, M. Tucholska, H. Choi, A.-C. Gingras and A. I. Nesvizhskii, Nature Methods, 2015, 12, 258–264.

49. D. B. Bekker-Jensen, O. M. Bernhardt, A. Hogrebe, A. Martinez-Val, L. Verbeke, T. Gandhi, C. D. Kelstrup, L. Reiter and J. V. Olsen, Nature Communications, 2020, 11, 787.

50. S. Keerthikumar, D. Chisanga, D. Ariyaratne, H. Al Saffar, S. Anand, K. Zhao, M. Samuel, M. Pathan, M. Jois, N. Chilamkurti, L. Gangoda and S. Mathivanan, Journal of Molecular Biology, 2016, 428, 688–692.

51. J. A. Fafián-Labora, J. A. Rodríguez-Navarro and A. O’Loghlen, Cell Metabolism, 2020, 32, 71-86.e75.

52. D. Muñoz-Espín and M. Serrano, Nature Reviews Molecular Cell Biology, 2014, 15, 482–496.

53. M. Borghesan, J. Fafián-Labora, O. Eleftheriadou, P. Carpintero-Fernández, M. Paez-Ribes, G. Vizcay-Barrena, A. Swisa, D. Kolodkin-Gal, P. Ximénez-Embún, R. Lowe, B. Martín-Martín, H. Peinado, J. Muñoz, R. A. Fleck, Y. Dor, I. Ben-Porath, A. Vossenkamper, D. Muñoz-Espin and A. O’Loghlen, Cell Reports, 2019, 27, 3956–3971 e3956.

54. E. M. Castaño, A. E. Roher, C. L. Esh, T. A. Kokjohn and T. Beach, Neurological Research, 2013, 28, 155–163.

55. T. Tanaka, A. Biancotto, R. Moaddel, A. Z. Moore, M. Gonzalez-Freire, M. A. Aon, J. Candia, P. Zhang, F. Cheung, G. Fantoni, R. D. Semba and L. Ferrucci, Aging Cell, 2018, 17, e12799.

56. J. P. de Magalhães, J. Curado and G. M. Church, Bioinformatics, 2009, 25, 875–881.

57. B. Lehallier, D. Gate, N. Schaum, T. Nanasi, S. E. Lee, H. Yousef, P. Moran Losada, D. Berdnik, A. Keller, J. Verghese, S. Sathyan, C. Franceschi, S. Milman, N. Barzilai and T. Wyss-Coray, Nature Medicine, 2019, 25, 1843–1850.

58. Z. Ahmed, H. Sheng, Y.-f. Xu, W.-L. Lin, A. E. Innes, J. Gass, X. Yu, H. Hou, S. Chiba, K. Yamanouchi, M. Leissring, L. Petrucelli, M. Nishihara, M. L. Hutton, E. McGowan, D. W. Dickson and J. Lewis, The American Journal of Pathology, 2010, 177, 311–324.

59. I. M. Riederer, M. Schiffrin, E. Kövari, C. Bouras and B. M. Riederer, Brain Research Bulletin, 2009, 80, 233–241.

60. C. López-Otín, M. A. Blasco, L. Partridge, M. Serrano and G. Kroemer, Cell, 2013, 153, 1194–1217.

61. I. Saez and D. Vilchez, Current Genomics, 2014, 15, 38–51.

62. Grillari, J., Hackl, M., Campisi, J., Kale, A., US Pat., WO2019002265A1, 2019.

63. D. Bhaumik, G. K. Scott, S. Schokrpur, C. K. Patil, A. V. Orjalo, F. Rodier, G. J. Lithgow and J. Campisi, Aging, 2009, 1, 402–411.

64. W.-F. Lai, M. Lin and W.-T. Wong, Trends in Molecular Medicine, 2019, 25, 673–684.

65. C. Cai, S. Min, B. Yan, W. Liu, X. Yang, L. Li, T. Wang and A. Jin, Aging, 2019, 11, 6371–6384.

66. M. Serpente, C. Fenoglio, M. D’Anca, M. Arcaro, F. Sorrentino, C. Visconte, A. Arighi, G. G. Fumagalli, L. Porretti, A. Cattaneo, M. Ciani, R. Zanardini, L. Benussi, R. Ghidoni, E. Scarpini and D. Galimberti, Cells, 2020, 9, 1443.

67. X. Li, M. Xu, L. Ding and J. Tang, Journal of Cancer, 2019, 10, 2836–2848.

68. Y. Yu and X.-C. Cao, Cancer Cell International, 2019, 19, 257.

69. K. S. Sheinerman and S. R. Umansky, Frontiers in Cellular Neuroscience, 2013, 7, 150.

70. K. S. Sheinerman, J. B. Toledo, V. G. Tsivinsky, D. Irwin, M. Grossman, D. Weintraub, H. I. Hurtig, A. Chen-Plotkin, D. A. Wolk, L. F. McCluskey, L. B. Elman, J. Q. Trojanowski and S. R. Umansky, Alzheimer′s Research & Therapy, 2017, 9, 89.

71. K. S. Sheinerman, V. G. Tsivinsky, L. Abdullah, F. Crawford and S. R. Umansky, Aging, 2013, 5, 925–938.

72. J. Muntel, T. Gandhi, L. Verbeke, O. M. Bernhardt, T. Treiber, R. Bruderer and L. Reiter, Molecular Omics, 2019, 15, 348–360.

73. L. Reiter, M. Claassen, S. P. Schrimpf, M. Jovanovic, A. Schmidt, J. M. Buhmann, M. O. Hengartner and R. Aebersold, Molecular & Cellular Proteomics, 2009, 8, 2405–2417.

74. L. Reiter, O. Rinner, P. Picotti, R. Hüttenhain, M. Beck, M.-Y. Brusniak, M. O. Hengartner and R. Aebersold, Nature Methods, 2011, 8, 430–435.

75. H. Wickham, Programming with ggplot2. In: ggplot2. Use R!, Springer, Cham, First edn., 2016.

76. M. Li, Y. Xia, Y. Gu, K. Zhang, Q. Lang, L. Chen, J. Guan, Z. Luo, H. Chen, Y. Li, Q. Li, X. Li, A.-a. Jiang, S. Shuai, J. Wang, Q. Zhu, X. Zhou, X. Gao and X. Li, PLoS ONE, 2010, 5, e11541.

