## Supplemental Figure 1 for "Comprehensive Profiling of Plasma Exosomes Using Data-Independent Acquisitions – New Tools for Aging Cohort Studies"

### Slide 1
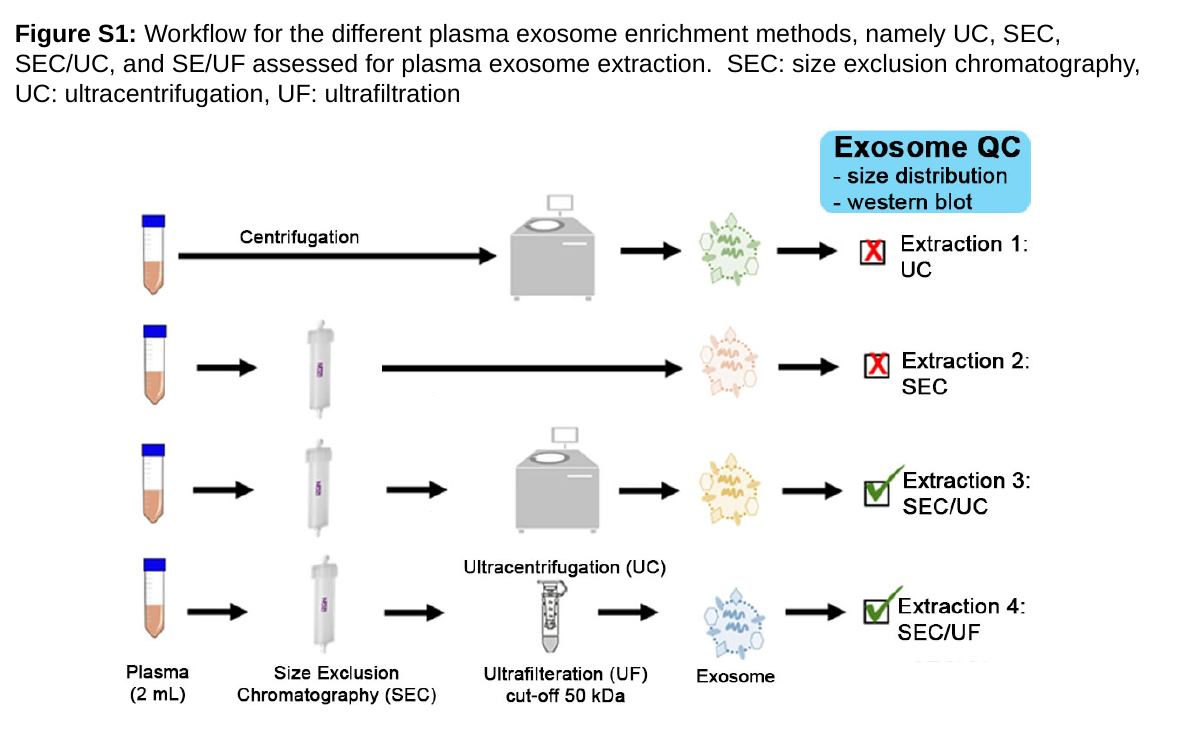

Figure S1: Workflow for the different plasma exosome enrichment methods, namely UC, SEC, SEC/UC, and SE/UF assessed for plasma exosome extraction. SEC: size exclusion chromatography, UC: ultracentrifugation, UF: ultrafiltration
