## Supplemental Figure 2 for "Comprehensive Profiling of Plasma Exosomes Using Data-Independent Acquisitions – New Tools for Aging Cohort Studies"

### Slide 1
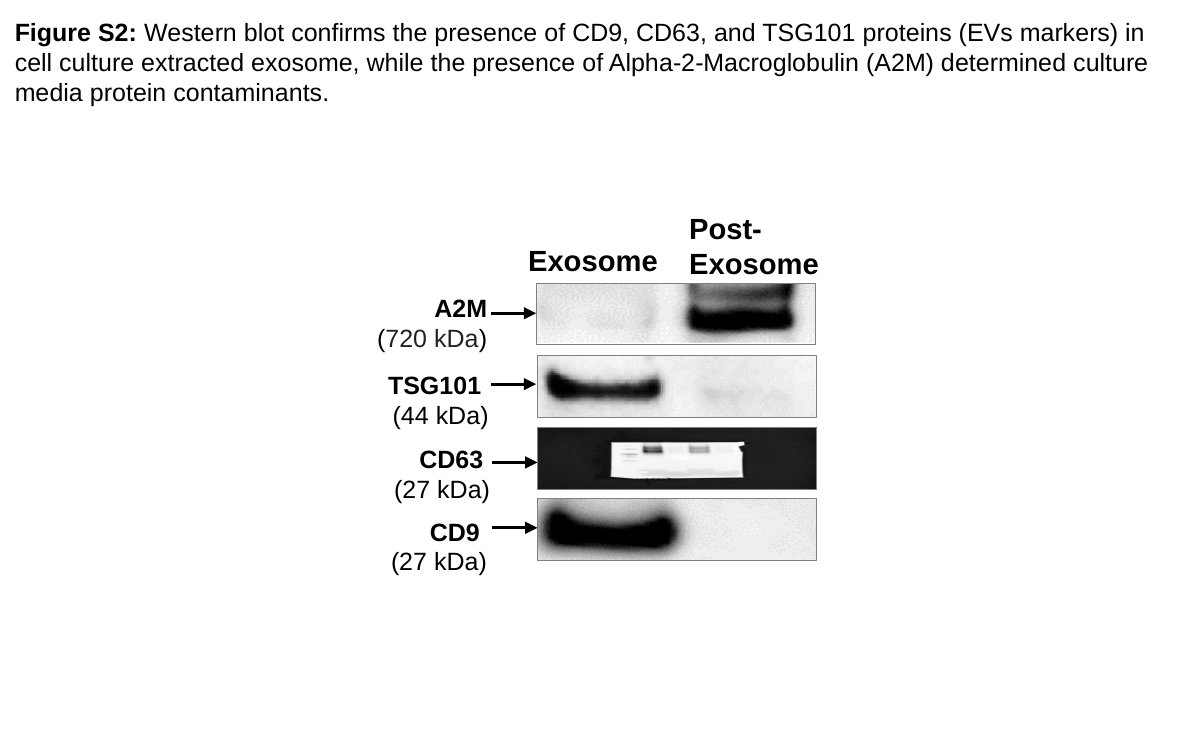

Figure S2: Western blot confirms the presence of CD9, CD63, and TSG101 proteins (EVs markers) in cell culture extracted exosome, while the presence of Alpha-2-Macroglobulin (A2M) determined culture media protein contaminants.
Post-Exosome
Exosome
TSG101
(44 kDa)
CD63
(27 kDa)
A2M
(720 kDa)
CD9
(27 kDa)
