## Supplemental Figure 3 for "Comprehensive Profiling of Plasma Exosomes Using Data-Independent Acquisitions – New Tools for Aging Cohort Studies"

### Slide 1
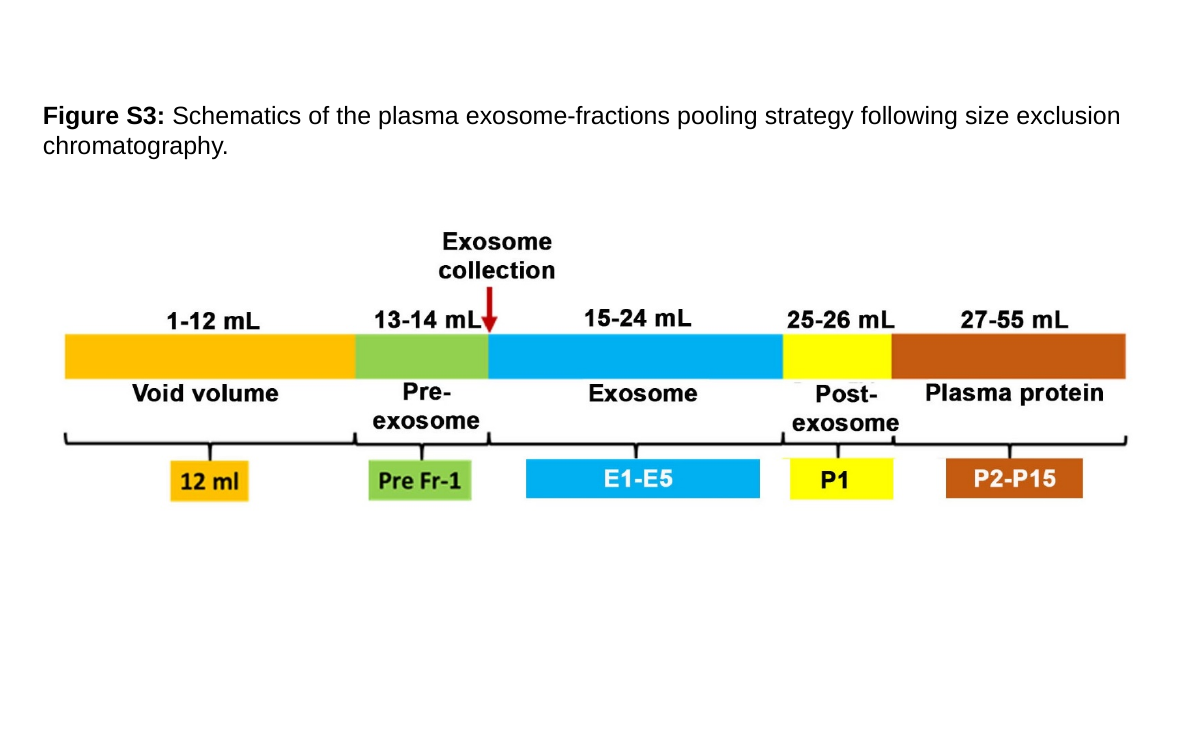

Figure S3: Schematics of the plasma exosome-fractions pooling strategy following size exclusion chromatography.
