## Supplemental Figure 4 for "Comprehensive Profiling of Plasma Exosomes Using Data-Independent Acquisitions – New Tools for Aging Cohort Studies"

### Slide 1
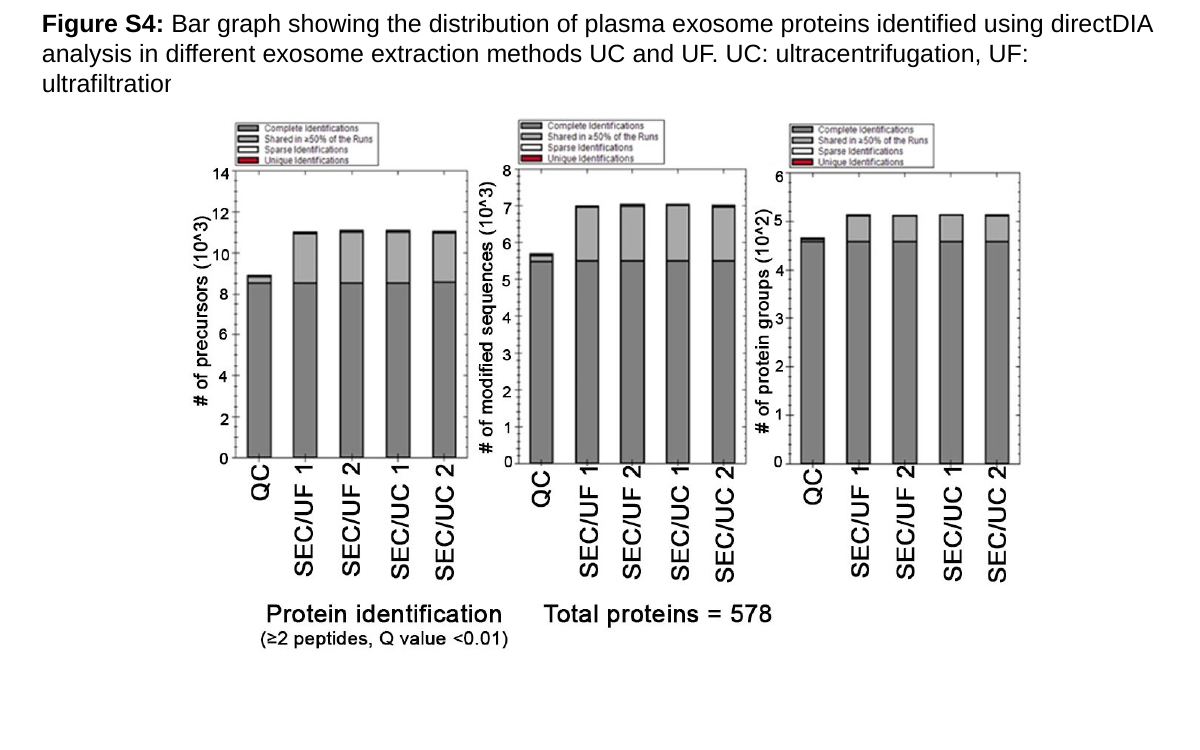

Figure S4: Bar graph showing the distribution of plasma exosome proteins identified using directDIA analysis in different exosome extraction methods UC and UF. UC: ultracentrifugation, UF: ultrafiltration
