## Supplemental Figure 5 for "Comprehensive Profiling of Plasma Exosomes Using Data-Independent Acquisitions – New Tools for Aging Cohort Studies"

### Slide 1
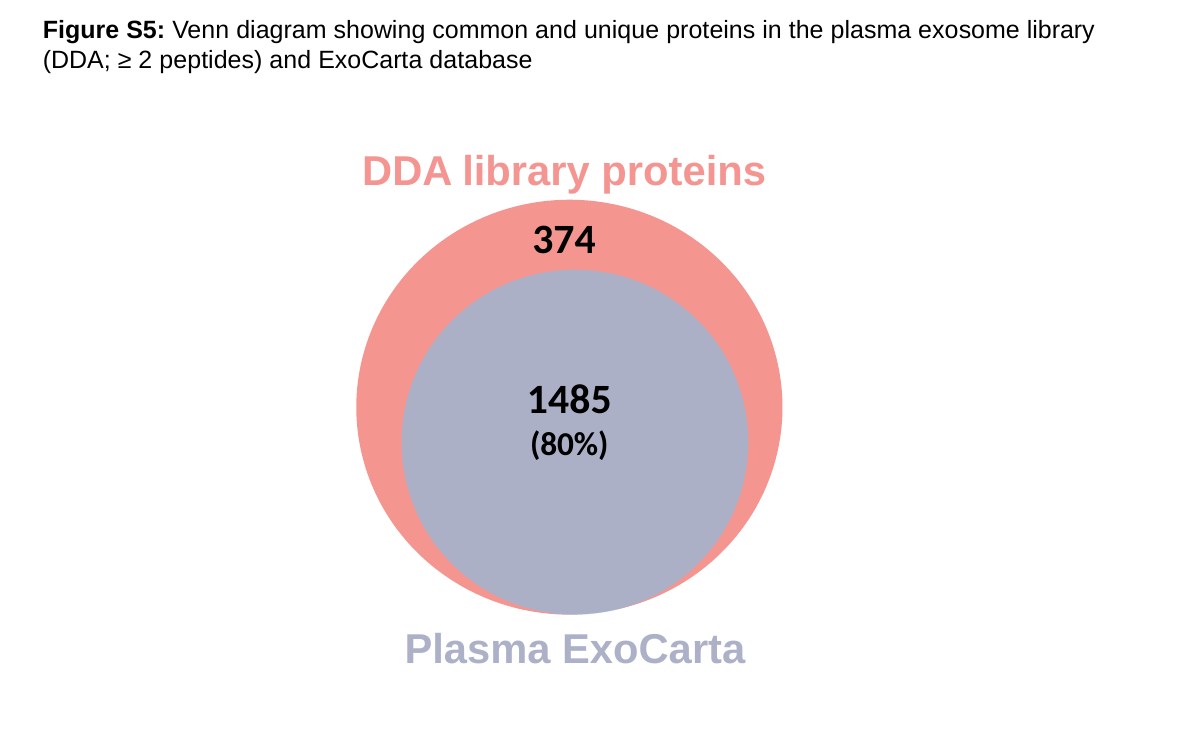

Figure S5: Venn diagram showing common and unique proteins in the plasma exosome library (DDA; ≥ 2 peptides) and ExoCarta database
DDA library proteins
Plasma ExoCarta
374
1485
(80%)
