## Supplemental Figure 6 for "Comprehensive Profiling of Plasma Exosomes Using Data-Independent Acquisitions – New Tools for Aging Cohort Studies"

### Slide 1
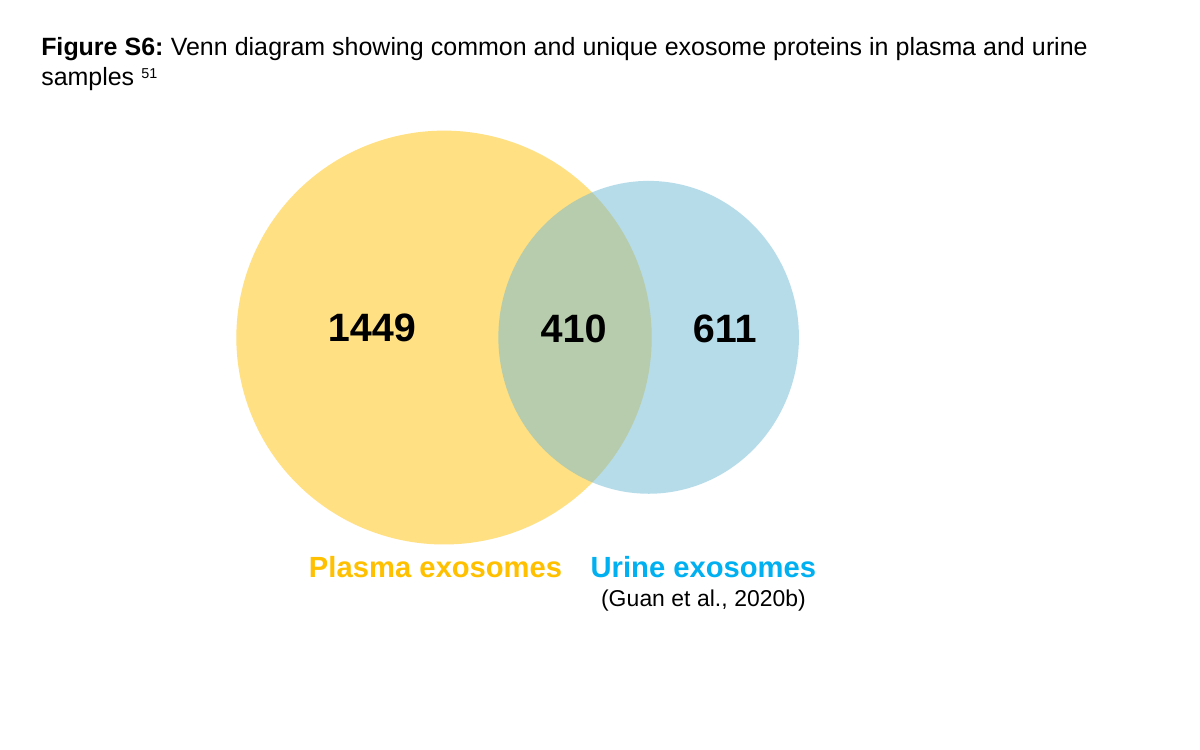

Figure S6: Venn diagram showing common and unique exosome proteins in plasma and urine samples ^51^
1449
410
Plasma exosomes
Urine exosomes
(Guan et al., 2020b)
611
