## Supplemental Figure 7 for "Comprehensive Profiling of Plasma Exosomes Using Data-Independent Acquisitions – New Tools for Aging Cohort Studies"

### Slide 1
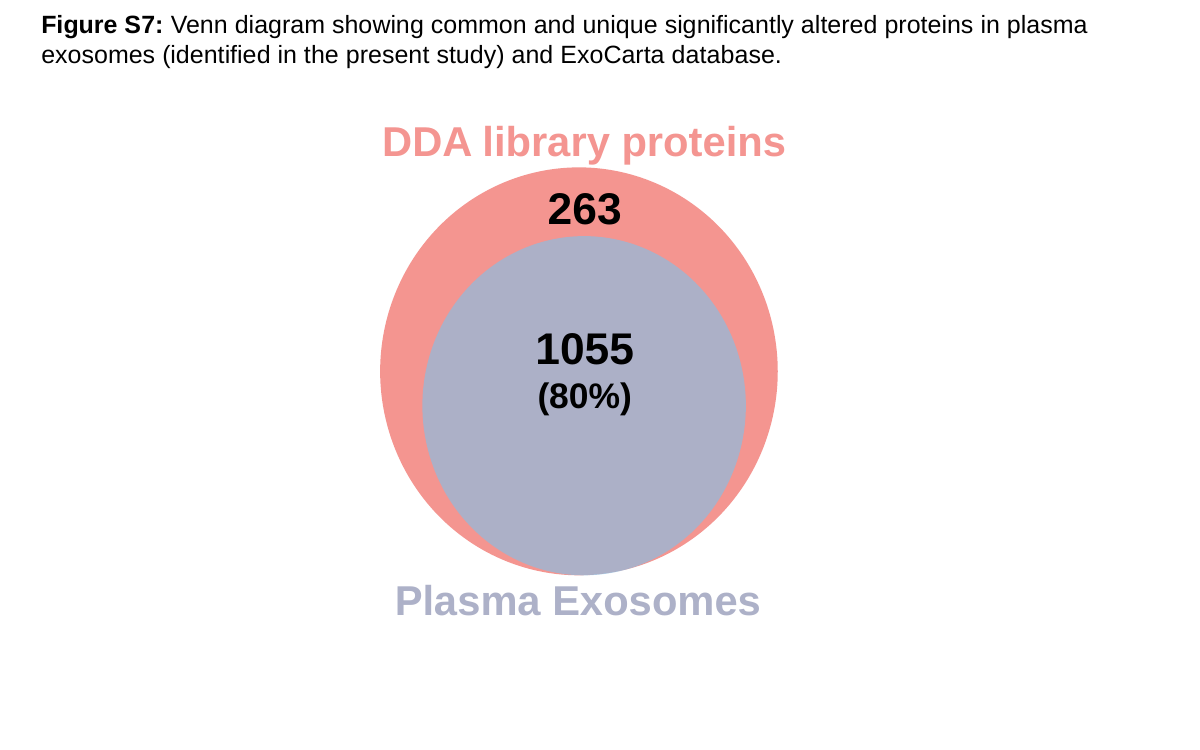

Figure S7: Venn diagram showing common and unique significantly altered proteins in plasma exosomes (identified in the present study) and ExoCarta database.
DDA library proteins
263
1055
(80%)
Plasma Exosomes
