## Supplemental Figure 8 for "Comprehensive Profiling of Plasma Exosomes Using Data-Independent Acquisitions – New Tools for Aging Cohort Studies"

### Slide 1
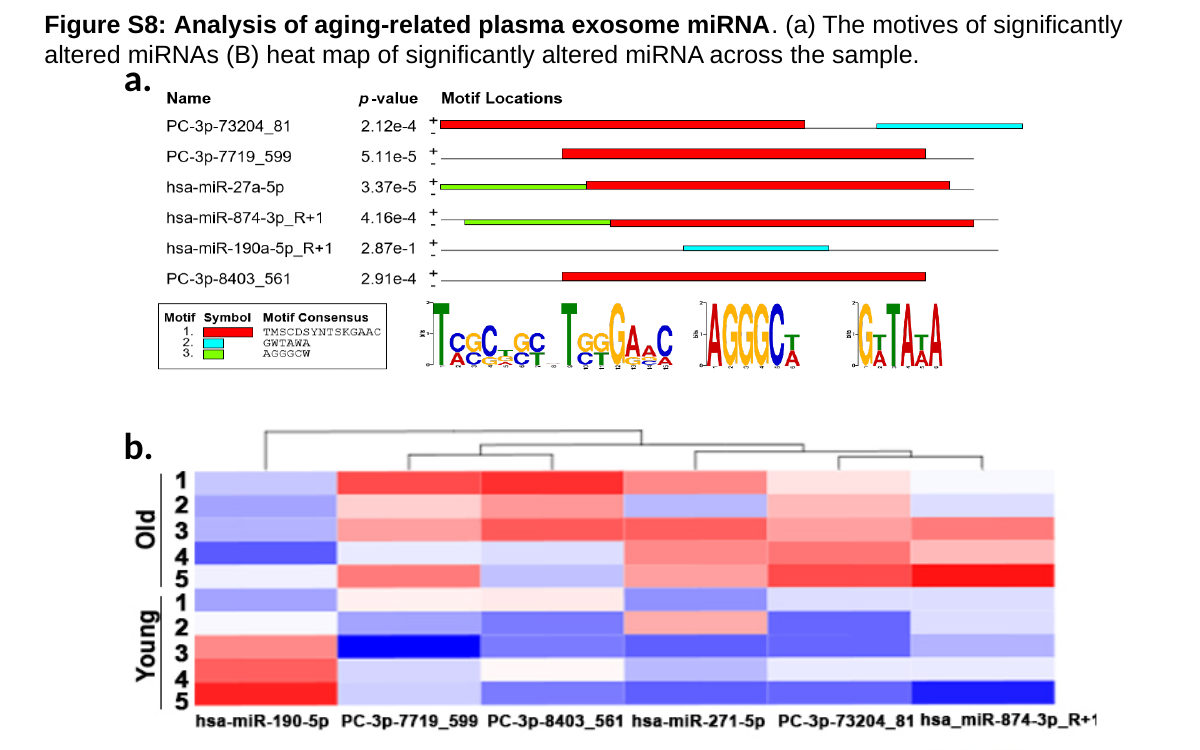

Figure S8: Analysis of aging-related plasma exosome miRNA. (a) The motives of significantly altered miRNAs (B) heat map of significantly altered miRNA across the sample.
a.
b.
